# NAD+ biosynthesis as a collateral lethality target for precision oncology

**DOI:** 10.1101/2022.12.13.518967

**Authors:** Florian L. Muller, Pingna Deng, Yu-Hsi Lin, Nikunj Satani, Naima Hammoudi, Gayatry Mohapatra, John M. Asara, Ronald A. DePinho

## Abstract

Genomic deletion of tumor suppressor genes (TSG) often encompasses neighboring genes which may be members of multi-gene families encoding cell essential functions. These genomic events create targetable cancer-specific vulnerabilities, termed “collateral lethality” as illustrated by homozygous deletion of the glycolytic gene *ENO1*, which sensitizes glioblastoma (GBM) cells to inhibition of its paralog *ENO2*. Here, we sought to generalize the concept by validating a second multi-gene family in an unrelated metabolic pathway. Nicotinamide-nucleotide adenylyltransferase (NMNAT) is an essential and (unlike the more extensively studied NAMPT) non-bypassable step in NAD biosynthesis encoded by three paralogs, one of which, *NMNAT1*, is homozygously deleted as part of the 1p36 tumor suppressor locus in TCGA data of GBM, Cholangiocarcinoma and Hepatocellular carcinoma (with near zero expression of found in several other malignancies). In a glioma cell line (Gli56) with homozygous deletion of *NMNAT1* expressing NMNAT2 and NMNAT3, shRNA-mediated knockdown of NMNAT2 is selectively toxic to Gli56 *NMNAT1*-deleted but not ectopically rescued cells. As NMNAT1 and NMNAT2 are predominantly localized in the nucleus and cytosol, respectively, these data suggest a functionally common pool of cytosolic and nuclear NAD+. Inducible shRNA-mediated extinction of NMNAT2 decreases NAD+ levels and selectively kills *NMNAT1*-deleted, but not *NMNAT1*-rescued cells *in vitro* and eradicates intracranial tumors *in vivo*. Thus, collateral lethality is a generalizable framework for the development of new classes of targeted agents with an informed clinical development path in cancer.

**Statement of significance:** An ongoing challenge in precision oncology is the translation of genomic data into actionable therapeutic opportunities with clear clinical benefit. We have demonstrated that genes homozygously deleted by virtue of chromosomal proximity to major tumor suppressor genes can confer cancer-specific vulnerabilities, termed “collateral lethality”. While our previous work validated one such collateral lethality target in glycolysis, we now provide empirical *in vitro* and *in vivo* evidence that this concept applies to another deleted gene governing an altogether distinct biochemical pathway. The generalization of collateral lethality may expand the spectrum of molecular targets in cancer with a genomically-informed path for accurate clinical development.

## INTRODUCTION

Homozygous genomic deletions centered on tumor suppressor genes are key drivers of tumor progression in diverse cancers (Cox et al., 2005, Proc Natl Acad Sci U S A). The stochastic nature of these genomic events frequently results in deletion of adjacent non-tumor suppressor genes, some of which play critical but redundant roles in normal cellular housekeeping. Our previous work has demonstrated that these passenger deleted genes can provide cancer-selective vulnerabilities, termed “collateral lethality” (Muller et al., 2012, Nature). This concept was validated in GBM which can sustain homozygous deletion of *ENO1* residing within the 1p36 tumor suppressor locus. ENO1 is a redundant member of a paralogous gene pair (ENO1/ENO2) playing a cell essential function in glycolysis. In this context, inhibition of the non-deleted paralogue ENO2 is profoundly toxic to cancer cells (with ENO1 deleted) but not normal cells which contain ENO1 activity. To assess whether collateral lethality serves as a general concept, we sought similar evidence for other cell essential paralogues resident in tumor suppressor loci.

The NAD+ biosynthesizing pathway has received increased attention with regards to development of chemotherapeutic drugs (Bi & Che, 2010, Cancer Biol Ther; Thakur et al., 2012, Biochem Biophys Res Commun). While precursors such as Nicotinamide and nicotinic acid readily permeate cells, nucleotides and NAD+ are unable to cross the plasma membrane, likely because of the multiple negatively charged phosphates (Nikiforov, Dolle, Niere, & Ziegler, 2011, J Biol Chem). The NAD+ biosynthesizing pathway is highly branched with multiple and distinct synthesis routes as salvage pathways. The only essential enzyme for both of the de-novo and salvage NAD+ biosynthesis is Nicotinamide-nucleotide adenylyltransferase (NMNAT). NMNAT is encoded by three, partially differential localized and expressed paralogues. Invertebrate data show that NMNAT is an essentially enzyme (Baba et al., 2006, Mol Syst Biol; Kim et al., 2010, Nat Biotechnol; Zhai et al., 2006, PLoS Biol), strongly suggesting that cancer cells with NMNAT1 deletion are dependent on the activity of either NMNAT2 or NMNAT3. NMNAT1 has the highest specific activity, is expressed ubiquitously and is localized predominantly to the nucleus while NMNAT2 is predominantly expressed in neural tissues associated with cytosolic face of endoplasmic reticulum/golgi membranes (Table S1). NMNAT3 is expressed ubiquitously but appears to be mitochondrial (Berger et al., 2005, J Biol Chem; Lau et al., 2009, Front Biosci (Landmark Ed)).

Here, we report that homozygous genomic deletions of the 1p36 locus in various cancers can include *NMNAT1*. Glioma cells lacking *NMNAT1* have exceptionally low NMNAT activity, which can be restored by ectopic re-expression of NMNAT1. ShRNA-mediated knockdown of NMNAT2 leads to depletion of NAD+ and profound disruptions in carbohydrate metabolism as well as to selective killing of *NMNAT1*-null, but not *NMNAT1*-rescued glioma cells. We further show that induction of inducible shNMNAT2 ablates intracranial NMNAT1-deleted intracranial orthotopic xenografted gliomas. These findings indicate redundant action of the two paralogues despite different subcellular localization, shedding new light on the NAD biosynthesis system and illustrate yet another exploitable collateral lethal vulnerability, highlighting its broad applicable framework for therapeutic development in oncology.

## RESULTS

### NMNAT1, an essential-redundant housekeeping gene, is homozygously deleted in cancer, including GBM

A review of the genomic status of the NAD-synthesizing pathway across diverse cancers (cBioportal.org, accessed 12/20/2013) indicates that the genomic alterations in most NAD-metabolism related genes (*NAMPT, NAPRT1, QPRT1, NADS*; Figure 1A) are amplifications or non-recurrent point mutations (Figure S1). The clear exception to this is NMNAT1 which resides in the 1p36 locus containing multiple tumor suppressor genes (Genovese et al., 2012, Cancer Discov; Henrich,Schwab, & Westermann, 2012, Cancer Res; Schlisio et al., 2008, Genes Dev;Wiedemeyer et al., 2008, Cancer Cell; Ying et al., 2010, Proc Natl Acad Sci U S A). This locus is frequently deleted in diverse cancers (Figure 1, S1 (Henrich et al., 2012,Cancer Res)) with homozygous deletion in 1-5% of GBM cases (Cancer Genome Atlas Research, 2008, Nature; Duncan et al., 2010, Oncotarget; Ichimura et al., 2008,Oncogene; Kotliarov et al., 2006, Cancer Res; Yin et al., 2009, Mol Cancer Res) and approximately 30% of these deletions encompass *NMNAT1* (Figure S1D). cBioPoral (https://www.cbioportal.org/) TCGA analysis of hepatocellular carcinoma (HCC or LIHC in TCGA code) indicates a similar fraction of *NMNAT1*-homozygous deletion cases (Figure S1D). Similar frequency of *NMNAT1* homozygous deletions is also observed in TCGA data for Cholangiocarcinoma (CHOL). Importantly – these deletion calls are confirmed in the ASCAT tumor-purity and whole-genome duplication correct Sanger Cosmic analysis (GRCh38 · COSMIC v97 https://cancer.sanger.ac.uk/cosmic/; Table S1). Patient-derived xenograft copy-number data from CrownBio (HuPrime®) also show presence of gastric adenocarcinoma (Model# GA2420). In addition to these *NMNAT1*-homozygously deleted tumors – many more tumors show near zero NMNAT1-expression due to a combination of heterozygous deletion and silencing by promoter methylation. We hypothesize that *NMNAT1* is likely a passenger event in the deletion or silencing of tumor suppressor genes are found both immediately upstream (*KIF1B* (Schlisio et al., 2008, Genes Dev)) and downstream (mir34a, *ERRFI1* (Genovese et al.,2012, Cancer Discov; Ying et al., 2010, Proc Natl Acad Sci U S A)) of its location.

**Figure 1:**
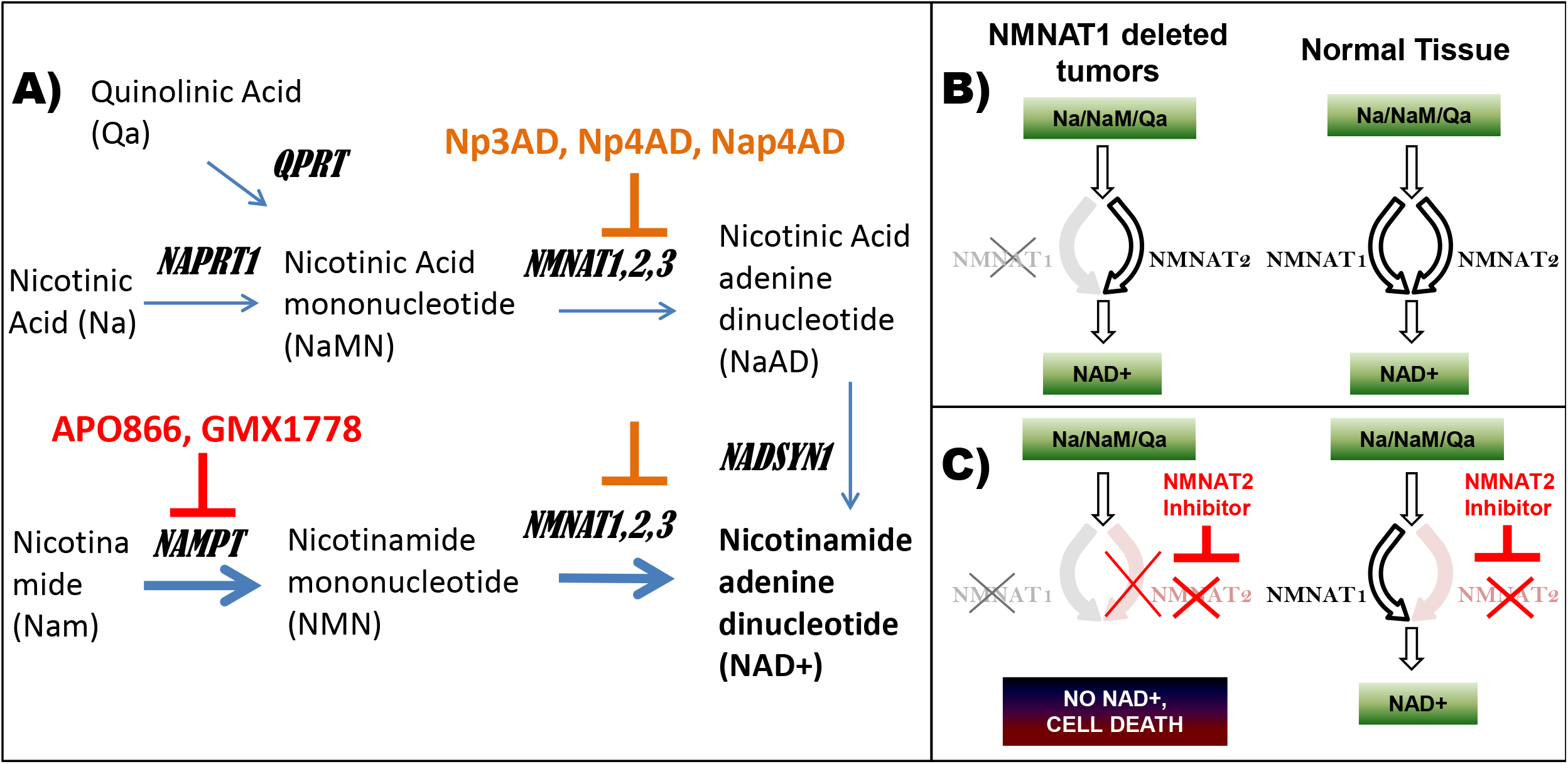
NMNAT2 as a collateral lethality target in *NMNAT1*-homozygously deleted tumors. NAD+ can be synthesized from three different precursors: quinolinate (requiring the *QPRT* gene, from tryptophan), nicotinate (*NAPRT*) and nicotinamide (*NAMPT*). Thickness of arrows indicates relative importance in flux levels in a typical cell, with nicotinamide pathway being the most important. Inhibitors of NAMPT currently in clinical trial are indicated in red. Tool compound inhibitors of NMNAT are indicated in orange (A). All NAD synthesis pathways ultimately require catalysis by NMNAT, which is encoded by three different genes. NMNAT3 is thought to localize to mitochondria and the mitochondrial NAD+ pool is thought to be segregated from that in the cytosol, hence, NMNAT1 and NMNAT3 are unlikely to be complementary. Therefore, NMNAT2 is likely responsible for cytosolic NAD synthesis in the absence of NMNAT1 (B). If that is indeed the case, tumors with 1p36 homozygous deletions covering *NMNAT1* should be highly sensitive to inhibition of NMNAT2, as this would fully eliminate cytosolic NAD synthesis (C).

A survey of our in house GBM cell line panel identified a *NMNAT1*-homozygously deleted GBM cell line, Gli56, derived from a recurrent GBM tumor (Figure 2A, (Mueller et al., 2007, Oncogene)). Triangulation with RNA-seq data confirmed absence of NMNAT1 mRNA in Gli56 cells (Figure 2B, C). In contrast, the D423-MG GBM cell line with 1p36 homozygous deletion spanning *ENO1* but not *NMNAT1*, shows expression of NMNAT1 (Figure 2B,C). RNA-seq data indicate exceptionally low levels of NMNAT1 mRNA in the NB1 neuroblastoma cell line, which was confirmed by Q-PCR (Figure S4C,D) and western blot (Figure 2D). The low levels of NMNAT1 in the NB1 cell lines are not the result of a genomic deletion but are typical of neuroectodermal-originating cells, which have very low levels of NMNAT1, but high levels of NMNAT2 expression (Figure S2). In contrast, GBM cell lines and astrocytes typically have very low levels of NMNAT2 mRNA (Figure S2, Figure S3) both in absolute terms and relative to the NB1 neuroblastoma cell line. Using whole cell lysates, NMNAT2 protein is readily detected by western blot in NB1 cells but is barely detectable in the above panel of GBM cell lines (Figure 2D). NMNAT2 is more readily detected in sub-cellular fractionated vesicular fractions, although it is still in much lower abundance in Gli56 than in NB1 cells (Figure S8A). RNA-seq data indicate that the NMNAT1-null Gli56 line shows a moderate elevation of NMNAT2 as compared to the other GBM cell lines and normal human astrocytes (Figure S4B). These results were confirmed by Q-PCR (Figure S4D).

**Figure 2:**
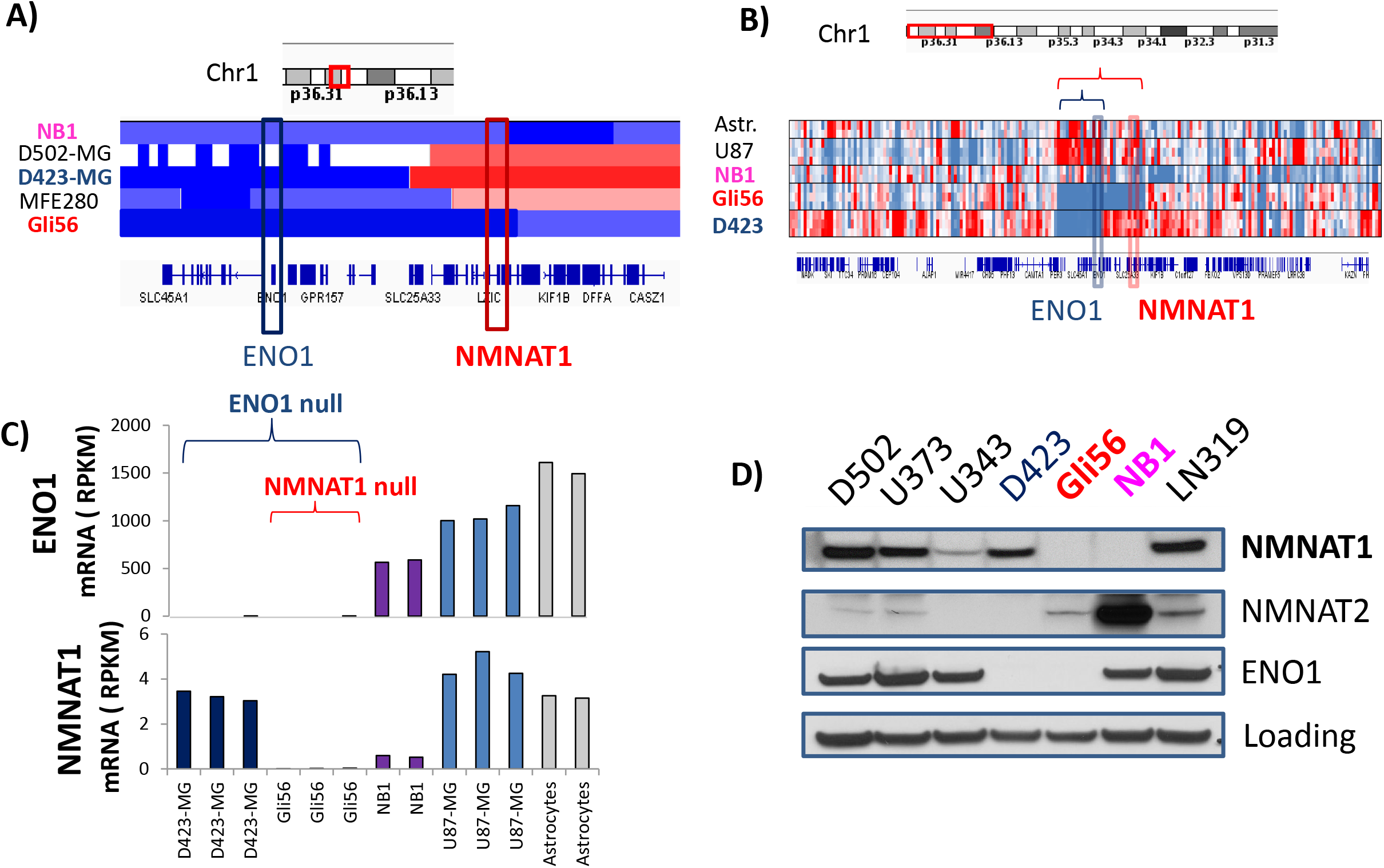
Homozygous deletion of NMNAT1 in the Gli56 glioma cell line. Gli56 is a cell line derived from a second resection of a recurrent GBM with a large 1p36 homozygous deletion covering ENO1 and NMNAT1 (**A**). The 1p36 homozygous deletion in NB1 (**A**) does not actually cover the *NMNAT1* gene (<u>Munirajan et al., 2008, J Biol Chem</u>). RNA-seq profiling data indicate close correlation between very low expression (blue) and genomic deletion (**B**) and confirm loss of ENO1 expression in both D423-MG and Gli56 cells, but loss of NMNAT1 expression restricted to Gli56 cells carrying the deletion (**C**). Western blot data confirm loss of ENO1 in both the Gli56 and D423-MG cell lines but loss of NMNAT1 protein expression in Gli56 only (**D**). The NB1/C201neuroblastoma cell line (<u>Munirajan et al., 2008, J Biol Chem</u>) shows scant NMNAT1 expression; this expression is low but not absent as the band becomes visible after prolonged exposure (data not shown).

Even so, despite this possibly compensatory increase, the level of total NMNAT enzymatic activity is dramatically lower in Gli56 *NMNAT1*-null cells compared to other cancer cell lines (Figure S5). In fact, the level of NMNAT activity is barely detectable above background in Gli56 *NMNAT1*-null cells (Figure S5), reflecting the fact that *NMNAT1* has the highest specific activity and accounts for the vast majority of NMNAT enzymatic activity in most cell types (Chiarugi et al., 2012, Nat Rev Cancer). Conversely, ectopic expression of *NMNAT1* restored NMNAT activity in Gli56 *NMNAT1*-deleted cells (Figure S8B, Figure S5).

### shRNA knockdown of NMNAT2 selectively depletes NAD+ and kills NMNAT1 deleted GBM cells in vitro

We posit that NMNAT2 is most likely to exert redundancy with NMNAT1 and sustain cellular NAD+ production in Gli56 cells. NMNAT3 is localized to the mitochondrial matrix, whereas NMNAT2 is localized to the outer face of cytosolic vesicles. As such the latter but not the former, shares the same pool of soluble metabolites as NMNAT1 (Chiarugi, Dolle, Felici, & Ziegler, 2012, Nat Rev Cancer). To validate NMNAT2 as collateral lethality target in *NMNAT1*-homozygous deleted gliomas, we performed stable shRNA knockdown using the pGIPZ vector against NMNAT2 as well as NMNAT3 in Gli56 *NMNAT1*-deleted, and an isogenic *NMNAT1*-rescued cells, where NMNAT1 is expressed ectopically. Knockdown of NMNAT3 was toxic to both rescued and NMNAT1 deleted cell lines (Figure S6B), underscoring the importance of mitochondrial NAD+ for cell viability, independently of cytosolic NAD+ pools. On the other hand, shRNA knockdown of NMNAT2 strongly inhibited the growth of Gli56 *NMNAT1*-null, but not rescued cells (Figure S6C), consistent with the notion that NMNAT2 sustains cytosolic NAD synthesis in the face of *NMNAT1*-deletion. Because of confounding factors with lentiviral infection delivered shRNA, we repeated this experiment using an isogenic system with doxycycline (DOX)-inducible shRNA against NMNAT2 in the pTRIPZ vector (using the same shRNA sequences, shNMNAT2-1 and 2) and in the pLKO vector (independent sequence, shNMNAT2-3). Efficiency of knockdown after DOX induction of shRNA was verified by Q-PCR (Figure S7A) and western blot (Figure 3B, S8A). Knockdown of NMNAT2 via either the pTRIPZ and pLKO vectors strongly and selectively killed *NMNAT1*-deleted but not rescued glioma cells (Figure 3A, Figure S7B; S8C, S11). Induction of shNMNAT2 resulted in selective apoptosis in *NMNAT1*-deleted glioma cells as evidenced by a dramatic upregulation of cleaved caspase 3 (Figure 4, Figure S12) and cleaved PARP (Figure 4). Again, this was completely prevented by ectopic re-expression of NMNAT1. Targeted tandem mass spectrometry (LC-MS/MS) measurements show that knockdown of NMNAT2 selectively decreases NAD+ levels (Figure 3C, S9B) in *NMNAT1*-deleted but not rescued glioma cells. This result was verified using a chromogenic photometric assay (Figure S10). NNMNAT2 knockdown also selectively decreased NADP+ levels, but much more modestly than NAD+ (Figure 3C, S9C).

**Figure 3:**
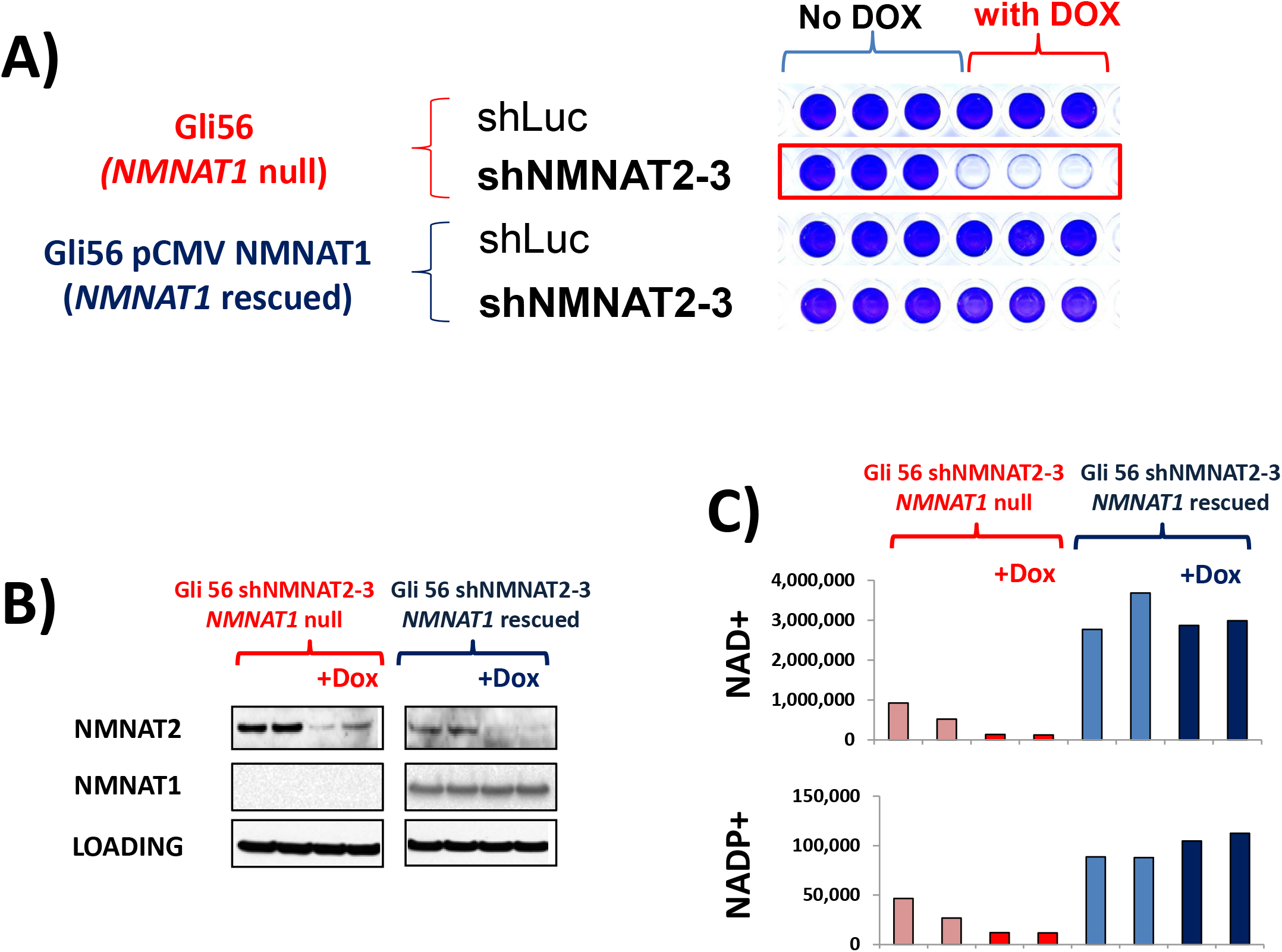
Selective depletion of NAD+ and toxicity in *NMNAT1*-deleted glioma cells by inducible shRNA against NMNAT2. Gli56 parental NMNAT1-deleted and Gli56 NMNAT1-rescued glioma cells were infected with pLKO DOX-inducible shRNA against NMNAT2 (shNMNAT2-3) or non-target (shLuciferase). Knockdown of NMNAT2 induced strong selective toxicity only in NMNAT1-deleted but not rescued Gli56 glioma cells (A). Knockdown of NMNAT2 following induction of the shRNA by DOX was confirmed by Western blot (B); Knockdown of NMNAT2 selective depleted NAD+, and to a lesser extent, NADP+, in NMNAT1 deleted but not rescued glioma cells (C). Data in Panel C are expressed in LC-MS/MS SRM peak areas.

**Figure 4:**
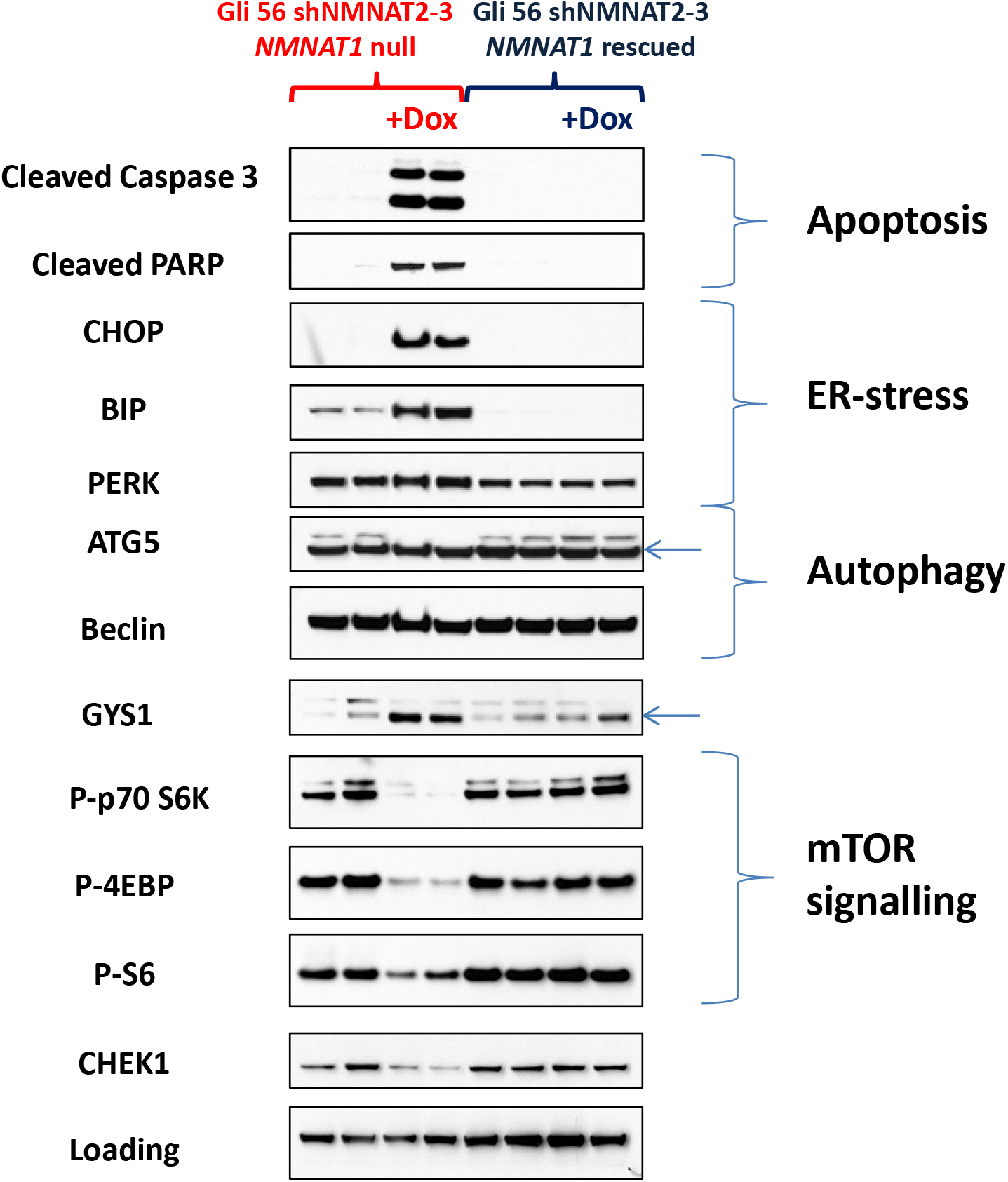
Knockdown of NMNAT2 leads selective induction of ER stress and induction of apoptosis. Proteins representative of pathways selectively altered by NMNAT2 knockdown in NMNAT1-deleted cells in the RPPA dataset (Suppl Table S1) were confirmed by western blot (red: *NMNAT1*-deleted, blue NMNAT1-rescued, shNMNAT2-3 induced by DOX). These include selective induction of apoptosis (Cleaved Caspase 3, Cleaved PARP), suppression of the mTOR signaling axis (p-S6, p-4EBP, p-p70 S6K), carbohydrate metabolism (GYS1), DNA damage sensing (CHEK1) and induction of ER-stress (BIP, CHOP).

We note that the effectiveness of cell killing by shNMNAT decreases with each culture passage even in the absence of DOX. Thus, after approximately 3 months of continuous passage post-lentiviral infection with shNMNAT2, DOX induction of the hairpin yielded only modest reductions in NMNAT2 mRNA and protein levels (<50%), as well as equally modest reductions in NAD+ levels. Consequently, the toxicity of shNMNAT2 induction was considerably attenuated (data not show). Promoter silencing of shRNA hairpins, even inducible ones, is a recurring technical problem, likely to be especially severe in this system, given that Gli56 is a hyper-methylating cell line (Mueller et al., 2007, Oncogene).

In *NMNAT1*-intact cancer cell lines, knockdown of NMNAT2 had minimal phenotypic effects, even in the low NMNAT1-expressing NB1 cell line (Figure S8C). Despite a 95%-knockdown of NMNAT2 in NB1 cells using the same pTRIPZ inducible hairpins (Figure S8A), there was only a slight decrease in NAD+ levels and cell growth was not materially affected (Figure S8C). Interestingly, even after a 95% knockdown, the absolute expression level of NMNAT2 was still higher in NB1 cells than in non-shRNA ablated Gli56 cells (Figure S8A).

Thus, taken together, these results indicate that sufficient NAD+ production can be sustained by surprisingly low levels of NMNAT activity.

### Depletion of NAD+ leads to profound carbohydrate metabolism remodeling, induction of ER stress and apoptosis

We wished to understand the biochemical mechanisms by which NAD+ depletion comprises cellular viability and leads to cell death. Inducible shRNA against NMNAT2 in *NMNAT1*-deleted and isogenic rescued glioma cells provides an ideal system to probe the biochemical consequences of NAD+ depletion. We performed unbiased proteomic (Reverse Phase Protein Array, RPPA, (Sheehan et al., 2005, Mol Cell Proteomics)) and metabolomic analysis (Polar compounds, (Yuan, Breitkopf, Yang, & Asara, 2012, Nat Protoc)) to understand the biochemical mechanisms by which of NAD+ depletion leads to cell death. Samples were collected after 4 days of induction of the shRNA (shNMNAT2-3) at a time when the bulk of *NMNAT1*-null, shNMNAT2-induced cells still appear morphologically normal, but scattered apoptotic cells begin to appear.

In *NMNAT1*-deleted but not *NMNAT1*-rescued glioma cells, metabolomic data revealed a profound accumulation of glycolytic intermediates upstream of Glyceraldehyde-3-phosphate dehydrogenase (GAPDH, fructose-6-phosphate,1,6-fructosebisphosphate, glyceraldehyde-3-phosphate) in response to induction of shNMNAT2, while at the same time, glycolytic intermediates downstream of GADPH are considerably depleted (e.g. 1,3-diphosphoglycerate, 3-phosphoglycerate, phosphonenolpyruvate; Figure 5, Table S2). It is noteworthy that GAPDH is the major NAD+ utilizing enzyme in glycolysis and these data suggest that this reaction becomes rate limiting with increasing depletion of NAD+ by the combined loss of *NMNAT1* through deletion and shRNA knockdown of NMNAT2. Consistent with this observation, metabolites around other key NAD(P)+ utilizing enzymes exhibit similar patterns. For example, induction of shNMNAT2 leads to a large accumulation of 6-phosphoglogluconate and its associated metabolites (gluconate, gluconolactonate), consistent with impairment of the NADP+ utilizing enzyme, 6-phosphogluconate dehydrogenase (PGD). Interestingly, RPPA data indicate upregulation of glucose-6-phosphate dehydrogenase (G6PD) as well as glycogen synthase (GYS1, Table S3, Figure 4), suggesting attempted compensation for depletion of NADPH in the former case and an attempt to sequester excess phosphor-hexoses in the latter. Indeed, RPPA data further support a stimulation of glycogen synthesis as the expression of GYS1 is not only increased but its phosphorylation (p-GYS1) is additionally decreased ((Zois,Favaro, & Harris, 2014, Biochem Pharmacol), phosphorylation is inhibitory to enzymatic activity, Table S3). This is likely to be the consequence of high levels of phosphorylated hexoses, which are known to stimulate glycogen synthesis (Zois et al., 2014, Biochem Pharmacol). Other particularly striking alterations include depletion of 3-phosphoserine, a metabolite downstream of the NAD+ utilizing enzymes GAPDH and phosphoglycerate dehydrogenase (PHPD). While such metabolite alterations can be directly linked to NAD(P)+ decreases, there are other alterations that defy easy explanation. Uracil and its associate metabolites are severely decreased (Uracil, UDP-glucose, UTP, UDP, N-carbamoyl-L-aspartate), although the uracil biosynthesis pathway is not strictly dependent on cytosolic NAD(P)+. Indeed, this depletion of uracil and UDP results in secondary disruptions, such as the hexose amine pathway, where the substrate of the UDP-dependent enzyme UDP-N-acetylglucosaminediphosphorylase (N-acetyl-glucosamine-1-phosphate) accumulates and the product (UDP-N-acetyl-glucosamine) is depleted in response to induction of shNMNAT2. These data indicate that the selective depletion of NAD+ by shNMNAT2 results in widespread metabolic disruptions, both directly linked to NAD(P)+ depletion as well as secondary to such effects. We suggest that such metabolite alterations may serve as useful target engagement markers for hypothetical therapeutic small molecule inhibitors or oligonucleotides targeting NMNAT2 in *NMNAT1*-deleted tumors.

**Figure 5:**
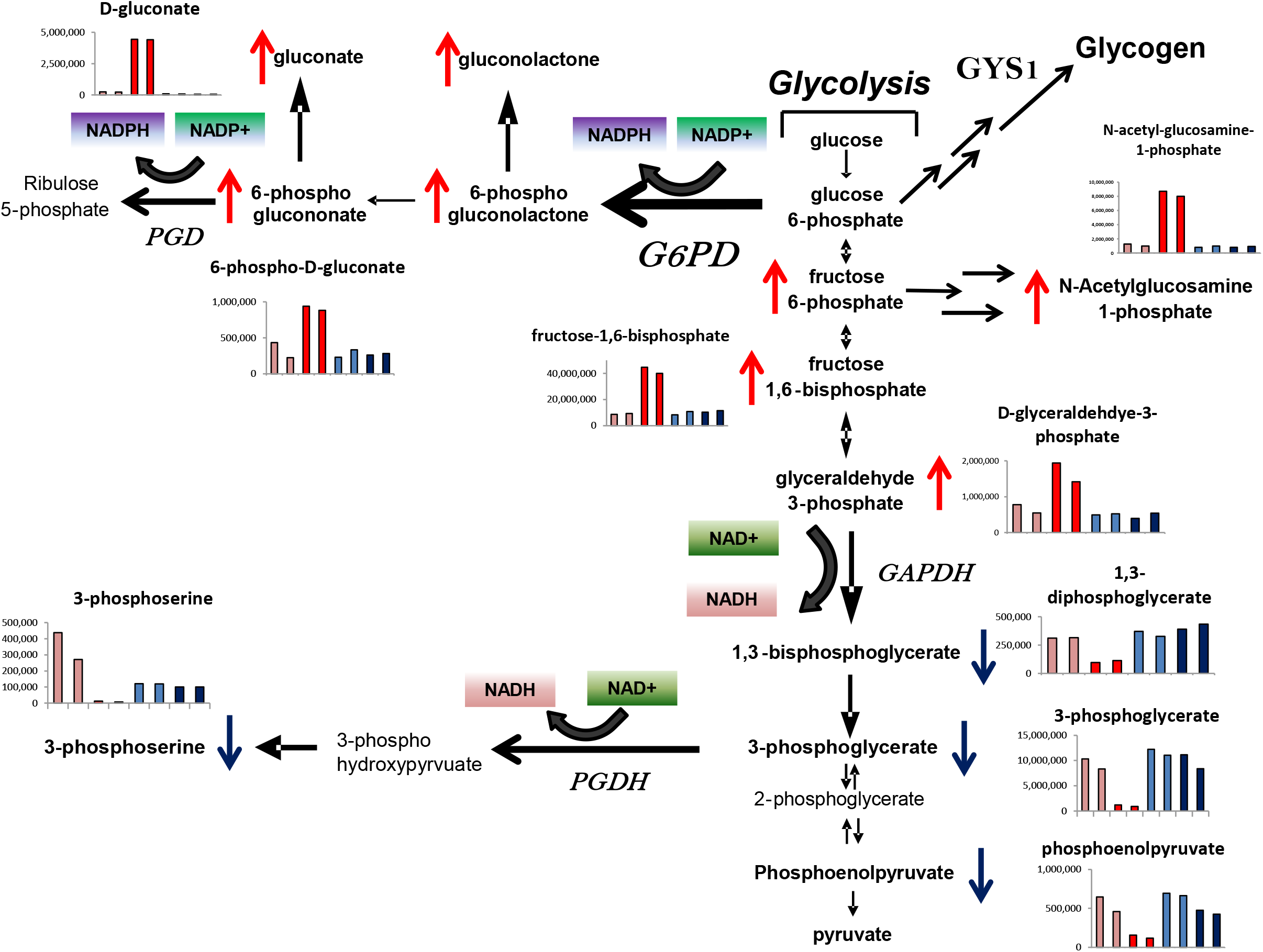
Knockdown of NMNAT2 leads to profound disruptions of NAD+/NADP+ dependent carbohydrate metabolism. Metabolites in the glycolysis and associated carbohydrate metabolism were mapped onto their respective position in the pathway, with selected metabolite alterations from Supplementary Table S2 being shown. Metabolites that were selectively increased by shNMNAT2 induction in *NMNAT1*-deleted cells are accompanied by a red arrow, while those showing a selective decrease are accompanied by a blue arrow. Induction of shNMNAT2-3 led to profound alterations in carbohydrate metabolites consistent with impairment of enzymatic steps requiring NAD(P)+ in Gli56 parental *NMNAT1*-deleted glioma cells (non-induced, light crimson bars, shNMNAT2 induced, red bars) but not NMNAT1-rescued (non-induced, light blue bars, shNMNAT2 induced, dark blue bars). This includes profound accumulation of metabolites upstream, and concomitant decrease in metabolites downstream, of the major NAD(P)+ utilizing enzymes (shown in italic, Glucose-6-phosphate dehydrogenase-G6PD; the enlarged arrow indicates RPPA data showing upregulation by induction of NMNAT2, 6-phosphogluconate dehydrogenase-PGD, Glyceraldehyde 3-phosphate dehydrogenase-GAPDH and Phosphoglycerate dehydrogenase-PGDH).

The consequences of NAD+ depletion had distinct effects on molecular signaling events associated with cell death, cell growth and cell proliferation. RPPA data confirmed by western blot indicate strong repression of cellular anabolic pathways, including a comprehensive decrease in the mTOR signaling pathway (dramatic decreases inp-S6, p-S6K, p-4EBP); at the same time, markers of cell proliferation presented a contradictory picture, with the phosphorylation of RB1 (initiation of G1/S) substantially reduced, but with the expression of the cell cycle regulated proteins, CDK1 and PCNA, largely unaltered (Table S3). This suggests an inability of the cell cycle regulatory machinery to properly sense the stress imposed by NAD+ depletion and downregulate cell proliferation; accordingly, this in turn likely derives from the multiple genomic hits affecting multiple tumor suppressor genes present in the recurrent glioma Gli56 cell line. At the same time, molecular signaling alterations provide strong evidence for induction of apoptosis. Cleaved Caspase 3, 7, as well as cleaved PARP are substantially and selectively upregulated in response to shNMNAT2 induction in NMNAT1 deleted but not rescued cells (Table S3 and Figure 4).

There is some evidence of attempted but evidently futile, compensation of depletion of NAD+. For example, there is an increase in G6PDH, which may indicate an attempt to upregulate NADPH generation. There is a coordinated upregulation of mitochondrial proteins (increased ATP5H, SDHA, PPIF) which may constitute an increase in mitochondrial biogenesis in an attempt to compensate for disrupted energy generation by glycolysis. Evidently, these attempts are ultimately futile, as induction of shNMNAT2 eventually kills *NMNAT1-* deleted glioma cells.

### Synergistic toxicities with NMNAT2 knockdown

To gain a better understanding of the mechanisms of toxicity and to obtain insight into *in vivo* biological consequences, we next queried what *in vitro* experimental conditions would cooperate with selective killing of NMNAT2 knockdown. We first asked our alterations in nutrients and co-factors would influence cell killing, by altering the media. We find that substitution of RPMI media (and to a lesser extent EMEM) instead of DMEM, dramatically accelerated selective killing by NMNAT2 knockdown (data not shown). DMEM is an extraordinarily rich medium, in which amino acids and vitamins (including nicotinamide) are present at four-fold higher levels than in RPMI. We hypothesized that increased cell killing by NMNAT2 knockdown in RPMI was due to decreased availability of nicotinamide in RPMI; we thus performed the same experiment in nicotinamide-free DMEM, however, this did not material affect response to shNMNAT2. We suggest that alternative production, via tryptophan catabolism is responsible for the bulk of NAD+ biosynthesis in Gli56 NMNAT1-null cells.

Because the glioma tumor microenvironment is characterized by severe hypoxia, which is known to modulate tumor response to therapy, we asked how hypoxia would affect toxicity of shNMNAT2 in *vitro*. Using a Tri-gas incubator set to 1% O2, we find that cell killing of NMNAT1 null Gli56 by shNMNAT2 is dramatically accelerated (Figure S6). We propose that this increased toxicity derives from the fact that NAD+ levels are significantly decreased under hypoxia (Figure S10) and/or that there is increased reliance on glycolysis which depends on NAD+ at the GAPDH step. Regardless of mechanism, it indicates that selective killing of glioma cells by selective ablation of NMNAT2 is more potent under condition prevalent in glioma *in vivo*.

We next asked whether NMNAT1 deletion alone or in combination with NMNAT2 knockdown, would sensitize cancer cells to known tool-compounds targeting the NAD+ system. Specifically, we asked whether inhibitors of NAMPT (FK866, (Bajrami et al., 2012, EMBO Mol Med)) or PARP (Olaparib, (Bajrami et al., 2012, EMBO Mol Med)) would synergize or additively kill with shNMNAT2 knockdown. FK866 was very toxic (nM IC50) but this did not materially differ between *NMNAT1*-null, rescued, with or without shNMNAT2 (data not shown).

### Inducible shRNA knockdown of NMNAT2 selectively eradicates NMNAT1 null intracranial orthotopic tumors

To test the potential *in vivo* anti-tumor effectiveness of NMNAT2 inhibition in the context of NMNAT1 deletion, we generated intracranial orthotopic tumors with Gli56 *NMNAT1*-null and *NMNAT1* rescued cells, carrying inducible shNMNAT2. One month after glioma cell injections, large established tumors became visible by T2-MRI (Figure 5 A,C,B for coronal and G,I,K for axial views). At this point, animals were treated with doxycycline supplied by both food and water to induce shNMNAT2. After 15 days of treatment, a second T2-MRI was taken (Figure 5 B,D,F and H,J,L, for coronal and axial views, respectively). For NMNAT1 null derived tumors, the induction of shNMNAT2 resulted in complete elimination of tumors in 3 out of 5 mice (compared panels C versus D and I versus K in Figure 5). These tumors have not recurred at the time of this writing (5 months since beginning of DOX treatment). In the two other animals, despite lack of complete regression, tumor growth was strongly suppressed. At the same time, induction of shNMNAT2 in *NMNAT1*-rescued tumor failed to suppress tumor growth (Figure 5 E versus F and K versus L) and led to the eventual death of all animals. Similar results were obtained with an independent shRNA against NMNAT2 (Figure S13).

## DISCUSSION

Cancer cells exhibit multiple metabolic alterations that have been explored as potential vulnerabilities for anti-cancer therapies, including addiction to glucose, glutamine and serine (Barthel et al., 1999, J Biol Chem; Cairns, Harris, McCracken, & Mak, 2011, Cold Spring Harb Symp Quant Biol; Galluzzi, Kepp, Heiden, & Kroemer, 2013, Nat Rev Drug Discov; Koppenol, Bounds, & Dang, 2011, Nat Rev Cancer; Levine & Puzio-Kuter, 2010, Science; Schulze & Harris, 2012, Nature; Tedeschi et al., 2013,Cell Death Dis; Wise et al., 2008, Proc Natl Acad Sci U S A). Cancer cells typically also exhibit a high turnover of NAD+ due to futile cycles of ADP-ribosylation (Petrelli,Felczak, & Cappellacci, 2011, Curr Med Chem; Stefano, Manerba, & Vettraino, 2013,Curr Top Med Chem; Vettraino, Manerba, Govoni, & Di Stefano, 2013, Anticancer Drugs). NAD+ is an essential co-factor in diverse biochemical reactions, including de-acetylation, ADP-ribosylation and in numerous oxidation/reduction reactions (Dolle, Skoge, Vanlinden, & Ziegler, 2013, Curr Top Med Chem). NAD+ is also the immediate precursor for NADP(H), thus also indirectly serving as a key intermediate for the generation of reducing equivalents required for biosynthetic and antioxidant biochemical reactions (Girolamo, Fabrizio, Scarpa, & Paola, 2013, Curr Top Med Chem; Petrelli et al., 2011, Curr Med Chem). As such, efforts have been made to target the NAD synthesis pathway for the treatment of cancer. The enzyme nicotinamidephosphoribosyltransferase (NAMPT), a key enzyme in the NAD+ salvage pathway (Figure 1A), has been investigated as a possible novel chemotherapeutic target in cancer (Christensen et al., 2013, J Med Chem; Shames et al., 2013, Clin Cancer Res). NAMPT inhibitors have entered phase I clinical trials, however, published studies suggest limited clinical success. Possibly, this may be due to the fact that NAMPT blockage can be bypassed by both the nicotinic acid salvage and the de-novo NAD biosynthesis pathway (Figure 1A). At the same time, it may be difficult to establish a therapeutic window given the importance of NAD+ in normal cellular metabolism.

We have previously proposed that a large therapeutic window can arise for a cancer drug targeting a cell-essential metabolic pathway, provided that such a drug targets the remain member of a pair of redundant paralogs where one is eliminated by genomic [passenger] deletions (Muller et al., 2012, Nature). The concept of collateral lethality (Muller et al., 2012, Nature) differs from general approaches to target cancer metabolism in that it seeks to use endogenous genetic differences between normal tissue and cancer to establish a therapeutic window. While we have previously experimentally validated one example of this strategy, whereby homozygous deletion of ENO1 on the 1p36 locus dramatically sensitizes to inhibition of ENO2, in order to demonstrate its general applicability, we sought to validate additional examples.

Looking no further than the immediate genomic neighborhood of ENO1, we identified homozygous deletion of the NAD+ biosynthesis gene, NMNAT1, as a candidate sensitizer for collateral lethality. NMNAT catalyzes the second to last reaction in NAD+ biosynthesis and is necessary for both the de novo and salvage pathways. Invertebrate data suggest that NMNAT enzymatic activity is most likely indispensable for cell viability: in species with only one NMNAT paralogue, it is an essential gene (*S. pombe, Drosophila, E. Coli* (Baba et al., 2006, Mol Syst Biol; Kim et al., 2010, Nat Biotechnol; Zhai et al., 2006, PLoS Biol)). In species with multiple homologues, individual isoforms are non-essential but the combined loss of all paralogues is lethal (*S. cerevisiae*,(Costanzo et al., 2010, Science; DeLuna et al., 2008, Nat Genet;Deshpande et al., 2013, Cancer Res)). Mammals have three NMNAT paralogues: *Nmnat1* and *Nmnat2* null mice are not viable at the organismal level (Conforti et al.,2011, FEBS J; Hicks et al., 2012, PLoS One), but neither *Nmnat1* nor *Nmnat2* are essential at the cellular level (Table S1). Nevertheless, heterozygotes and even sub-heterozygote *Nmnat2* mice with <25% of normal levels, are viable (Gilley, Adalbert, Yu, & Coleman, 2013, J Neurosci) and *Nmnat2*^-/-^*Sarm1*^-/-^ are fully viable – indicating that a major mechanism of toxicity in *Nmnat2* null mice may be activation of Sarm1 rather than depletion of NAD+ per se. Rather surprisingly *NMNAT3* is non-essential in human cancer cell lines (DepMap) and *Nmnat3*^-/-^ mice are viable with only minor defect in vertebrates (Table S1). NMNAT1, 2, 3 have distinct and overlapping expression patterns (Figure S2, S3A, Table S1 (Berger, Lau, Dahlmann, & Ziegler, 2005, J Biol Chem; Lau, Niere, & Ziegler, 2009, Front Biosci (Landmark Ed))). NMNAT1,2,3 catalyze the second to last step of NAD+ biosynthesis and have both unique and complementary roles.

TCGA CNV data indicate that *NMNAT1* is homozygously deleted in GBM as well as LIHC and CHOL as part of the 1p36 tumor suppressor locus. Congruent with this we identified a GBM cell line with a 1p36 deletion spanning *NMNAT1* and generated an isogenic rescued clone ectopically expressing NMNAT1. Knockdown of NMNAT2 by two independent sets of shRNA resulted in selective toxicity to *NMNAT1*-deleted but not isogenic rescued glioma cells. Visualized by T2-MRI, we observed complete eradication of intracranial tumors by NMNAT2 knockdown, which in the majority of cases did not recur after discontinuing shRNA knockdown by withholding doxycycline. This effect can be directly attributed to the lack of NMNAT1, as this sensitivity could be completely reversed by ectopic re-expression of NMNAT1. Biochemical profiling confirmed that as expected, shRNA knockdown of NMANT2 selectively decreased NAD+ and to a lesser extent NADPH+ in *NMNAT1* deleted cells. NMNAT isozymes share only about 30% identity and differ substantially in quaternary structure (NMNAT1 is a hexamer, while NMNAT2 is a dimer); NMNAT1 has the highest specific activity and is thought to be the major isoform in the cell. Indeed, even without induction of shRNA against NMNAT2, Gli56 NMNAT1 deleted cells had >90% lower overall NMNAT activity compared to NMNAT1-intact or NMNAT1-rescued cells. As such, the levels of NAD+ in Gli56 NMNAT1 deleted cells are sustained by comparatively low levels of NMNAT2.

Overall, these results recapitulate, expand and generalize what we had previously demonstrated with ENO1/ENO2. There are several striking parallels between the NMNAT1/2 and ENO1/2 collateral lethality pairs. ENO1 and NMNAT1 are homozygously deleted in GBM and are expressed broadly and constitutively. Deletion of each enzyme results in a >90% decrease in total enzymatic activity, be it ENO1 or NMNAT1. ENO2 and NMNAT2 would be the drug targets in these deleted tumors and their expression is restricted predominantly to the nervous system (Figure S3) as well as being significantly lower than that of NMNAT1. Indeed, in both cases, the metabolic pathway (glycolysis and NAD+ biosynthesis) is carried out by less than 10% of the normal enzymatic level. Given the much higher expression of NMNAT2 in the surrounding brain as compared to glioma cells, we suggest that relatively small levels of a NMNAT2 inhibitor would be sufficient to inhibit NAD synthesis in *NMNAT1*-deleted GBM tumors to reach toxic threshold. One critical difference between the ENO1/ENO2 and NMNAT1/NMNAT2 pairs is that while ENO2 null mice are viable, NMNAT2 null mice die shortly after birth, apparently due to axonal degeneration (Table S1). It is therefore conceivable that a NMNAT2 inhibitor might be too toxic to be tolerated. There are several reasons that lead us to believe this will not be the case. Mice with less than 25% of normal NMNAT2 levels are phenotypically normal (Table S1), suggesting that a 75% inhibition of NMNAT2 with a pharmacological inhibitor should be readily tolerated. Finally, *Nmnat2 Sarm1* double knockout mice are viable (Table S1) which suggests *Nmnat2* knockout drives axonal degeneration through deleterious activation of Sarm1 rather than depletion of NAD+ depletion.

Given the many fold higher expression of NMNAT2 in the surrounding brain as compared to *NMNAT1*-deleted glioma cells (Figure S3), we suggest that relatively small levels of a NMNAT2 inhibitor would be enough to inhibit NAD synthesis sufficiently in NMNAT1-null GBM tumors to reach toxic threshold. Indeed, a partial validation of this idea is provided by shRNA in the NB1 neuroblastoma cell line: even though a >90% knockdown of NMNAT2 is achieved by inducible shNMNAT2 (Figure S8A), the residual level of NMNAT2 is still sufficient to sustain near normal NAD+ generation (Figure S9) and maintain cell viability (Figure S8C).

Because NMNAT1, 2, & 3 differ substantially in sequence and structure (isozymes share ~30% identity and NMANT1 is a hexamer while NMNAT2 is a dimer), there is little reason to believe that NMNAT2 specific inhibitors could not be synthesized. Given that NMNAT1 is the major isoform of the enzyme in GBM cells, it may be unnecessary to develop a NMNAT2 specific inhibitor: a pan-NMNAT inhibitor may be specifically toxic to NMNAT1 homozygously deleted cells, as only low amounts of inhibitor would be required to eliminate the residual NMNAT activity sustaining NAD+ synthesis. It has been reported that gallotannin, a high molecular weight tanic acid polymer, can act as a general inhibitor of NMNAT activity with a *Ki* in the mid μM range (Berger et al., 2005, J Biol Chem). However, many other biological activities have been described for gallotannin (Xu, Du, Meiser, & Jacob, 2012, Nat Prod Commun), so its usefulness as a specific NMNAT inhibitor is likely limited, especially *in vivo*. A series of NAD analogues has been generated, some of which show potent and selective inhibition of NMNAT2 *in vitro* (Petrelli et al., 2011, Curr Med Chem). However, at this stage, these compounds are in the high μM range (Lauredana Cappelaci, personal communications).

While this work has focused on NMNAT2 as a collateral lethality target in *NMNAT1*-homozygously deleted tumors – our results do not rule out that NMNAT3 could also be an attractive opportunity – especially given that NMNAT3 appears to be dispensable both at the organismal and the cell-line specific level (Table S1).

On a final note, although the percentage of patients with NMNAT1 homozygous deletion is small, the therapeutic window for NMNAT2 inhibitors is likely to be very large, suggesting those patients will likely experience an exceptionally strong response. Furthermore, the number of GBM patients carrying NMNAT1 homozygous deletions may be considerably higher in recurrent as compared to primary tumors. The vast majority of genomic profiling studies, including the GBM TCGA, have focused on primary rather than recurrent tumor. In the case of glioblastoma, recurrence almost invariably occurs 12 months or so after primary tumor resection. Small-scale studies with paired primary/recurrent GBM indicate a ~3 fold increase in 1p36 deletions upon recurrence (Hulsebos, Troost, & Leenstra, 2004, J Neurol Neurosurg Psychiatry;Martinez, Rohde, & Schackert, 2010, J Neurooncol); though specific data on *NMNAT1* were not generated. In this light, it is noteworthy that the Gli56 cell line was derived from a case of recurrent rather than primary glioblastoma. Taken this into account, we speculate that the number of fatal cases of GBM carrying *NMNAT1*-homozygous deletions, hence patients benefiting from NMNAT inhibitors, may be considerably higher than the frequency estimated from the TCGA data (Figure S1).

We and others have hypothesized that collateral deleted genes may expose targetable vulnerabilities in cancer. Our present work experimentally demonstrates the redundant action of NMNAT2 in the generation of NAD+ in cancers cells with deletions encompassing *NMNAT1*. Taken together, these results suggest that these types of collateral lethality pairs are likely a common occurrence in the cancer genome and that the concept of collateral lethality may be a widely applicable therapeutic strategy.

**Figure 6:**
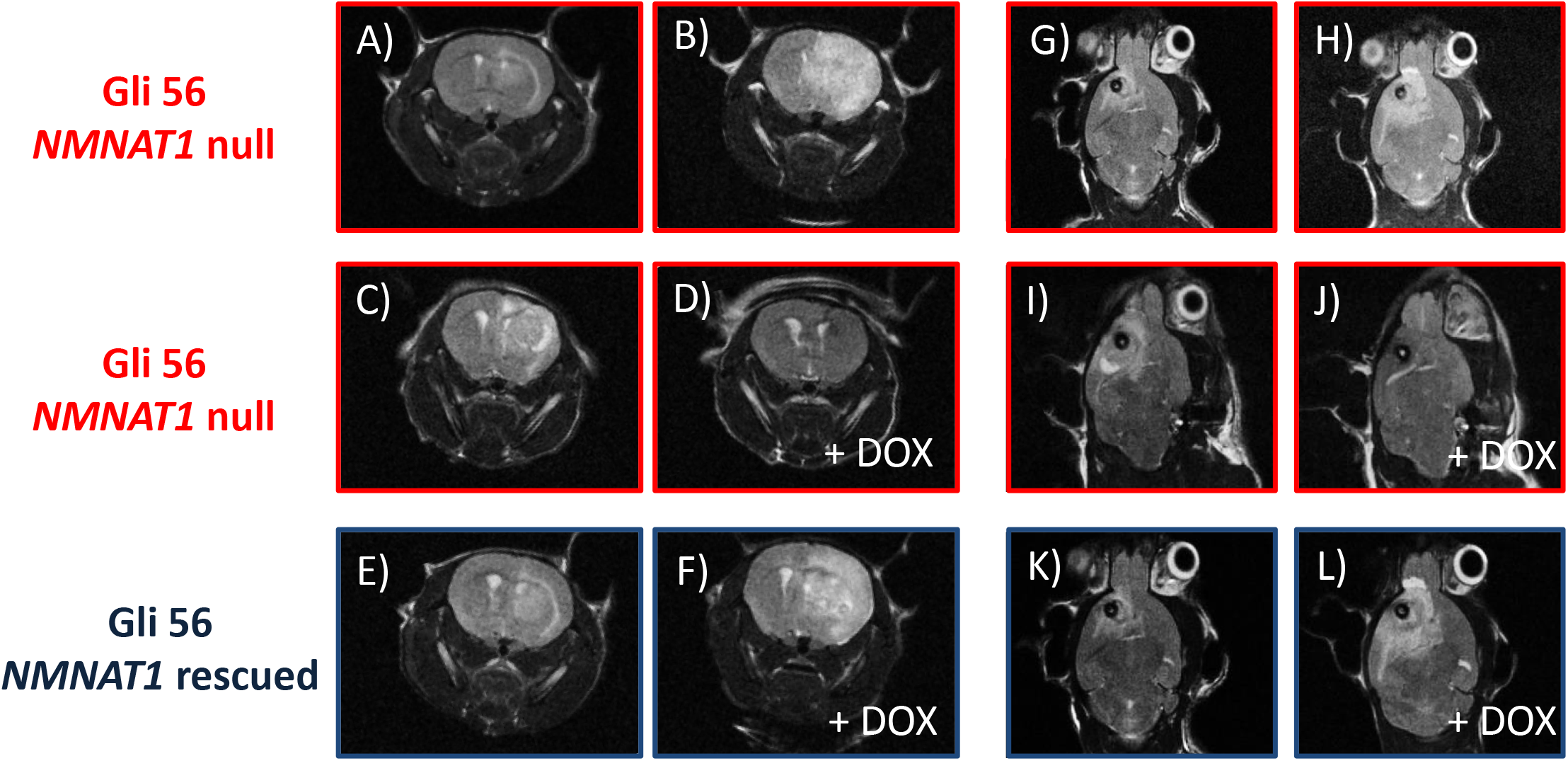
Inducible shRNA knockdown of NMNAT2 selectively eradicates NMNAT1-null intracranial gliomas. Coronal (A-F) and axial (G-L) T2-MRI images of three individual intracranial tumor bearing mice (each row is the same individual), taken 1 month (A,C,E,G,I,K) after intracranial cell injection and again 15 days later (B,D,F,H,J,L), during which time two mice (D,J and F,L) received DOX. Tumors are apparent by their white T2-hyperintensities. Three individual animals are shown: two bearing Gli56 *NMNAT1*-null tumors, one untreated (first row, A,B,G,H) and one receiving DOX to induce shNMNAT2 (second row, C,D,I,J); the third animal bearing Gli56 NMNAT1-rescued tumors (E,F,K,L) and received DOX (F,L). Non-dox treated *NMNAT1*-null tumors grew robustly over 15 days (A versus B, and G versus H), while dox-shNMNAT2 induction caused complete tumor regression (C versus D coronal, and I versus J, axial). Induction of shNMNAT2 does not shrink or attenuate growth in NMNAT1-rescued tumors (E versus F and K versus L).

## SUPPLEMENTARY MATERIALS LEGENDS

**Table S1: Expression and knockout phenotypes of mammalian NMNAT paralogs**

**Table S2: Polar compound profile of the effects of inducible NMNAT2 knockdown (shNMNAT2-3) in *NMNAT1*-deleted and *NMNAT1*-rescued Gli56 glioma cells.**

The table shows the metabolomic output, sorted as a fold change of the effects of NMNAT2 knockdown in *NMNAT1*-null cells. Experimental groups in columns are color-coded as follows: Gli56 *NMNAT1*-deleted shNMNAT2-3 without induction of the hairpin, light crimson; while those with induction of the shRNA with dox are coded in red. Gli56 *NMNAT1*-rescued shNMNAT2-3 without induction of the hairpin coded in light blue; while those with induction of the shRNA with dox are coded dark blue.

**Table S3: RPPA profile of the effects of inducible NMNAT2 knockdown (shNMNAT2-3) in *NMNAT1*-deleted and *NMNAT1*-rescued Gli56 glioma cells.**

Gli56 *NMNAT1*-deleted shNMNAT2-3 (without induction of the hairpin, light crimson; Four days after induction of the shRNA with dox, red). Gli56 *NMNAT1*-rescued shNMNAT2-3 (without induction of the hairpin, light blue; Four days after induction of the shRNA with dox, dark blue).

## METHODS

### Cell Culture

The cell lineD423-MG (1p36 homozygously deleted, excluding *NMNAT1*) kindly provided by Dr. Bigner (Duncan et al., 2010, Oncotarget). Gli56 (1p36 homozygously deleted, excluding *NMNAT1*) was obtained from David N. Louis (Mueller et al., 2007,Oncogene). The deletion in D423-MG spans the *CAMTA1, VAMP3, PER3, UTS2, TNFRSF9, PARK7, ERRFI1, SLC45A1, RERE, ENO1, CA6, SLC2A5, GPR157, MIR34A, H6PD, SPSB1*, and *SLC25A33* genes while the deletion in Gli56 spans the *UTS2, TNFRSF9, PARK7, ERRFI1, SLC45A1, RERE, ENO1, CA6, SLC2A5, GPR157, MIR34A, H6PD, SPSB1, SLC25A33, TMEM201, C1orf200, PIK3CD, CLSTN1, CTNNBIP1, LZIC, NMNAT1, RBP7* and *UBE4B* loci. The NB1/C201 (1p36 homozygously deleted, excluding *NMNAT1*) cell line was initially generated by Ohira *et al* (Ohira et al., 2000, Oncogene) and kindly shared by S. Schlisio (Karolinska Institute, Stockholm, Sweden). There are two distinct neuroblastoma cell lines named NB1 in the literature: NB1/C201 used here is identical to NB1 in the Sanger Cosmic dataset (Forbes et al., 2008, Curr Protoc Hum Genet) but not in Cancer Cell Line Encyclopedia dataset (Barretina et al., 2012, Nature). The 1p36 homozygous deletion in NB1 includes *KIF1B, PGD and UBE4B*. Cells were routinely cultured in high glucose, glutamine and pyruvate containing Dulbecco’s modified Eagle’s medium (DMEM) with 20% fetal bovine serum (FBS). For selected experiments, Eagle’s modified medium (EMEM) and Roswell Park Memorial Institute (RPMI) 1640 with glutamax were also used as indicated. U87-MG and SW1088 glioma cell lines were cultured as described previously. Normal human astrocytes were obtained from ScienCell. Nicotinamide (niacinamide) free media was obtained from D. Yoder (BIO5 Media Facility, U. Arizona, Tuscon, AZ). Nicotinamide and NMN were purchased from Sigma and added as indicated. For hypoxia experiments, a tri-gas incubator maintained at 1% O2 was employed (Thermo Scientific, Heracell 150i, 1.0% O2, 5% CO2).

### shRNA knockdown of NMNAT

shRNA knockdown was performed essentially as described previously (Muller et al.,2012, Nature). We screened 13 hairpins targeting NMNAT2 and found 2 independent sequences that reduced mRNA levels by >50%. Initial hairpin screenings were performed in the Expression Arrest pGIPZshRNAmir vectors (Thermo Scientific/Open Biosystems), while actual experiments were performed in the doxycycline inducible pTRIPZ vector (Thermo Scientific/Open Biosystems) using the sequences from the GIPZ screening. ThepTRIPZ*NMNAT2*shRNA sequences were as follows: V2LHS_100026(TTCCTTATGGCTCTCCAAG) and V3LHS_400732(TTATTTCTAAAGGAAACCT); shNMNAT2-1 and −2. These two hairpins were combined for all further inducible shRNA experiments. The same experiments were carried out for NMNAT3 V3THS_405269 (GGAAAGAATCAAGGAGG), V3HTS_400730 (GGAAAGAATCAAGGAGG). A non-targeting shRNA against firefly luciferase (shFF) was used as a control. For independent validation we used pLKO shRNAs in a doxycycline inducible vector. We also screen 5 pLKO hairpins for NMNAT2 with one shRNA showing >70% expression knockdown. The sequence was as follows: shpLKO NMNAT2-TRCN0000035439(TAGCTCAATCTCTTCATACCG), termed shNMNAT2-3.The shRNA expressing vectorswereintroduced into cancer cell lines by lentiviral infection. Recombinant lentiviral particles were produced by transient transfection of 293T cells following a standard protocol. Briefly, 30 μg of the shRNA plasmid, 19.5μg of pCMV Δ8.9, and 19.5 μg of psPAX2 were transfected using polyethylenimine (1μg/μl, Polysciences # 23966-2) into 293T cells plated in 6-well plates. Viral supernatant was collected 72h after transfection, centrifuged to remove any 293T cells and filtered (0.45μm). For transduction, viral solutions were added to cell culture medium containing 4 μg/mL polybrene; 48 h after infection, cells were selected using 2 μg/mL puromycin and tested for NMNAT2 knockdown by Q-PCR/western blotting.

### Ectopic expression of NMNAT1 in *NMNAT1*homozygously deleted cell lines

Rescue of the phenotypic effects of knocking down NMNAT2 in the cell line Gli56 was performed by re-expression of NMNAT1 from an ectopic plasmid. For the ectopic re-expression of NMNAT1, sequenced verified cDNA (Orfeome) clones were gateway cloned into the pHAGE-CMV lentiviral vector and lentivirally transduced into glioma cell lines as described above. Successful re-expression of NMNAT1 was verified by western blot.

### Proliferation assays

Cell growth of shRNA-treated cell lines was assayed either through crystal violet staining or using the Promega CellTiter-Glo proliferation kit (Roche) or alternatively by live-cell confluence measurements with the IncuCyte (Essen BioScience). Growth curves using the IncuCyte were generated by confluence imaging every 2 hours with quadruplicate replicates with and initial seeding of 10^3^ cells/well in 96-well plates. At the indicated time point (usually 2 weeks), cells were fixed with 10% formalin and stained with crystal violet solution for 1 h. Dye extraction was performed using 10% acetic acid solution, and absorbance was read at 595 nm. CellTiter-Glo experiments were performed according to the manufacturer’s instructions; 10^3^ cells/well were plated in a 96-well plate for each time point, and luminescence readings were taken every 24 h. All experiments were performed in triplicate.

### Apoptosis Assay

Apoptotic cells were counted by staining with YO-PRO®-1 Iodide (491/509, Life Technologies, Y3603). Apoptotic cells become permeable to YO-PRO®-1 while live cells are not stained. Total cell number was quantified by Hoechst 33342 (Cat# H3570 Invitrogen). Cells were seeded as 2×10^3^ cells/well in 96-well plates and treated in the presence or absence of doxycycline. On the day of apoptosis survey, old media was removed and each well was filled to 100 μl with fresh media. Hoechst and YO-PRO®-1 were mixed in 1:100 PBS and 10 μl of the mixture was added in each well gently without touching the cells for a final dilution of 1:1000 from stock. Plates were then incubated at 37°C for 2 hours. Image capture and quantification was done by High Content Screening System-Operetta (Perkin Elmer).

### Western blotting

Cells were washed twice in ice-cold phosphate-buffered saline (PBS), and incubated in RIPA buffer for 15 min with gentle rocking. Lysates were then collected, sonicated, and centrifuged at 14,000 rpm for 60 min at 4 °C. Protein concentration was measured using the BCA kit (Thermo Scientific - Pierce #23225). SDS-PAGE and western blotting were performed as described previously (Muller et al., 2012, Nature). The following antibodies were used: NMNAT2 (Abcam 56980, Sigma WH0023057M1); ENO1, NMNAT1 (Proteintech 11399-1-AP, Abcam # ab45548) and GAPDH CST# 3683; phosphor-AMPK Thr172 CST# 2535 from Cell Signaling Technologies and Vinculin from Sigma-Aldrich.Apoptosis was measured by western blots for Cleaved Caspase 3 (CST# 9664) and Cleaved PARP (CST#5625), BIP (CST# 3177), CHOP (CST# 2895), PERK (CST# 5683), ATG5 (CST# 12994) and Beclin 1 (CST# 3495). Other antibodies used were Glycogen Synthase 1 (Abcam# ab40810), Phospho-S6 (CST# 5364), Phospho-p70 S6 Kinase (CST# 9205), Phospho-4EBP1 (CST# 9451) and CHEK1 (Epitomics# 1250).

### Subcellular Fractionation

Subcellular protein fractionation kit for culture cells from Thermo Scientific (78840) was used to separate protein to cytosol, cytoplasmic member, and nuclear soluble compartment. The protocol from the manufacturer was followed to the letter. Cells were harvested using trypsin-EDTA, washed with PBS, and transferred at 2×10^6^ cells to a 1.5ml microcentrifuge tube. Briefly, cytosol was first separated out using a non-denaturing cell permeabilizing reagent, while membrane fractions and nucleous were separated using detergents of increasing strength. Protein concentration was determined using a BCA (Pierce™ BCA Protein Assay Kit #23225); protein concentrations were normalized, mixed with 4× loading buffer, the mixture was denatured and SDS-PAGE was performed as for western bloting. NMNAT2 (Abcam 56980, Sigma WH0023057M1), GAPDH (CST# 3683), Laminin A (Abcam, ab14055), and HSP19 (Uniprot P07901) antibodies were used.

### Transcriptomic profiling by RNA-seq and Q-PCR

RNA was isolated from ~10^6^ cells using Triazol extraction followed by purification with the QiagenRNAeasy kit as done previously (Paik et al., 2009, Cell Stem Cell; Sahin et al., 2011, Nature). RNA-seq was performed by the Sequencing and Microarray Facility (SMF) core at U.T.M.D. Anderson (Houston, TX). Libraries were generated using Illumina’sTruSeq kit and were sequenced using the Illumina HiSeq2000 Sequencer.Raw read RNA-seq data were mapped to hg19 reference genome using Bowtie (Langmead & Salzberg, 2012, Nat Methods). The mapped reads were then assembled by Cufflinks (Trapnell et al., 2012, Nat Protoc)to generate a transcriptome assembly for each sample. After the assembly phase, Cufflinks quantified expression level of the transcriptome in each gene for each sample (i.e., FPKM, fragments per kilobase of transcript per million fragments mapped).

For RT qPCR, RNA samples were reverse transcribed into cDNA using the High-Capacity cDNA Reverse Transcript kit (Life Technologies). cDNA samples were subjected to qRT-PCR quantification in duplicate and performed with Power SYBR Green PCR Master Mix (Invitrogen) according to the product guides on a Agilent Mx3005P machine.

The primer sequences used for real-time qRT-PCR are the following:

NMNAT1_9_A 5’-CCTTGCTTGTGGTTCATTCA-3’

NMNAT1_9_B 5’-TGCCTTTGACAACTGTGTA-3’

NMNAT2F1: 5’-CACCTGTGATCGGACAGCC-3’

NMNAT2R1: 5’-GTGCCCAGATTGGCATTCTC-3’

NMNAT2F2: 5’-ACGGTGATGCGGTATGAAGAG-3’

NMNAT2R2: 5’-CACCTCCATATCTGCCTCGTT-3’

NMNAT3_43_A 5’-CATGGCAAGGCACTCTTC-3’

NMNAT3_43_B 5’-TTGGGGGTCTGGAAGGTC-3’

Loading control

RPL39-F: 5’-CAGCTTCCCTCCTCTTCCTT-3’

RPL39-R: 5’-GCCAGGAATCGCTTAATCC-3’

### Metabolomics

Small-molecule metabolites from conditioned media, washed cells or tumors will be extracted with 80% methanol at −80°C (Yuan et al., 2012, Nat Protoc), following our published methodology (Ying et al., 2012, Cell). Methanol extraction recovers polar compounds such as most carboxylic acids, alcohols, sugars but not lipids. After extraction and centrifugation, the sample is lyophilized by freeze drying. Lyophilized samples were analyzed using microcapillary liquid chromatography tandem mass spectrometry (LC-MS/MS) using selected reaction monitoring (SRM) with positive/negative polarity switching on a hybrid 5500 QTRAP mass spectrometer (AB/SCIEX). 300 SRM transitions target >260 polar metabolites (Yuan et al., 2012, Nat Protoc) which includes most major glycolytic and citric acid cycle intermediates and importantly, most of the elements in the NAD/NADH biosynthesis pathway. MS-peak quantification was performed using MultiQuant 2.1 software and relative levels of each metabolite in Q3 peak area units across samples.

### Chromogenic spectrophotometric NAD+ assay

For routine NAD+ quantification, a spectrophotometric, enzyme-linked NAD+ assay was used (Enzychrom™, Bioassay systems, Hayward, CA). The assay measures the oxidation of MTT (yellow) to formazan (blue), the rate of which is dependent on the levels of NAD+. Cell extracts were prepared according to the manufacturers’ instructions and normalized by protein concentration. Spectrophotometric absorption at 565nm was quantified using a Pherastar plate reader (BMG Lab system). For samples with low NAD+ levels, read times were increased beyond the 30 minutes specified in the kit.After spectroscopic quantification, a standard curve established with an NAD+ standard provided with the kit was used to express NAD+ in absolute levels, which was then normalized by protein concentration.

### NMNAT enzymatic activity assay

NMNAT enzymatic activity was quantified in native, non-denatured lysates of cancer cell lines following the method of Orsomando *et al* (Orsomando et al., 2012, PLoS One), with slight modifications. Native lysates were prepared with 20mM Tris (pH 7.4), 1mM EDTA, 1mM mercaptoethanol (Muller et al., 2012, Nature). After scraping, cells were broken by soniation and lysates cleared by centrifugation at 15,000 rpm for 15 minutes.

Protein concentration was determined and normalized across samples. The reference assay mixture (0.4 mL final volume) contained 100mMTriethanolamine, pH 7.5, 25 mM MgCl2, 20 mM NaF, 1 mM both NMN and ATP, and 100 μg of cell lysate. In this assay, NMN and ATP are converted to NAD+, which was left to run for 30 minutes; after this incubation, the assay was stopped and NAD+ was measured using the spectrophotometric chromogenic assay described in the section above. Controls included samples with lysis buffer only and reaction buffer which omitted NMN (substrate of NMNAT) as well as inhibition by 100μM Gallotanin (Sigma T0200).

### Orthotopic brain tumor formation

Intracranial cell injections were performed by the M.D. Anderson Intracranial Injection Core at M.D. Anderson (Dr. Fred Lang, Director, (Lal et al., 2000, J Neurosurg)). Intracranial tumors were established by injection of 400,000 cells into the brain of immunocompromised female mice (Taconic), aged 6-9 weeks. The animals were first bolted (a bolt is a plastic screw with a hole in the middle which driven into the skull, (Lal et al., 2000, J Neurosurg)), allowed to recover for 2 weeks, and glioma cells were injected through the bolt with a Hamilton syringe. Bolting and cell injections were performed by the M.D. Anderson Intracranial injection fee-for-service core (Lal et al.,2000, J Neurosurg). Animals were sacrificed for humane reasons when neurological symptoms became apparent. Doxycycline was provided to the animals as indicated, in the form of Dox Diet (Protein 20.8%, Fat 8.7%, Fiber 2.1%, Ash 4.8%, Moisture < 10%, Carbohydrate 53.6%, Calories 3.75 kcal/gm), which contains 200mg of doxycycline per kg of diet (BioServ, S3888) and water (Dox 2mg/ml, Sucrose 40mg/ml). All procedures were approved by the M.D. Anderson Animal Care and Use Committee.

### MRI evaluation of intracranial tumor growth and regression

MRI measurements were performed in a 4.7 T Biospec USR47/40 (Bruker Biospin MRI, Billerica, MA) at the M.D. Anderson small animal imaging facility (SAIF). Briefly, animals were maintained under deep anesthesia with isoflurane with body temperature maintained by a heating blanket. Anesthetized mice were restrained with the head held in a stereotactic holder. Breathing was monitored and synchronized with the instrument. Routine tumor detection was done by T2-weighed imaging. First, a low resolution axial scan is taken to center the field properly, after which a series of high resolution axial and coronal scans are recorded. Gli56-derived gliomas exhibit very strong T2-MRI signals, including early diffuse hyperintensities, and during later stages, extensive edema.

## ACKNOWLEDGMENTS

We thank Susanne Schlisio (Karolinska Inst) for sharing NB1 cells, Verlene Henry for technical assistance with intracranial cell injections, C. Kingsley, K. Michael and J. Bankson for excellent technical support for MRI imaging. We thank Trang Tieu and Tim Heffernan for help with preparation of the pTRIPZ and inducible pLKO shRNAs and Simona Colla for help with design of NMNAT1 rescued cells. We thank M. Ziegler for useful discussion on the biology of NMNAT and primer sequences for NMNAT paralogues. We thank Marina Protopopova for help with high imager experiments proliferation and apoptosis experiments. We thank Per Wu for RNA seq analysis. We thank Joseph R. Marszalek and Y. Alan Wang for key discussions and suggestions. We thank Min Yuan and Susanne Breitkopf for help with the mass spectrometry experiments. This work was funded by NIH-NCI grants 7P01CA095616-10 (R.A.D), 5P01CA120964 (J.M.A.), 5P30CA006516 (J.M.A.), CPRIT RP140612 (R.A.D), NIH SPORE (F.L.M.).

## REFERENCES

Baba, T., Ara, T., Hasegawa, M., Takai, Y., Okumura, Y., Baba, M., Mori, H. (2006). Construction of Escherichia coli K-12 in-frame, single-gene knockout mutants: the Keio collection. Mol Syst Biol, 2, 2006 0008. doi: 10.1038/msb4100050

Bajrami, I., Kigozi, A., Van Weverwijk, A., Brough, R., Frankum, J., Lord, C. J., & Ashworth, A. (2012). Synthetic lethality of PARP and NAMPT inhibition in triple-negative breast cancer cells. EMBO Mol Med, 4(10), 1087–1096. doi: 10.1002/emmm.201201250

Barretina, J., Caponigro, G., Stransky, N., Venkatesan, K., Margolin, A. A., Kim, S., … Garraway, L. A. (2012). The Cancer Cell Line Encyclopedia enables predictive modelling of anticancer drug sensitivity. Nature, 483(7391), 603–607. doi: 10.1038/nature11003

Barthel, A., Okino, S. T., Liao, J., Nakatani, K., Li, J., Whitlock, J. P., Jr., & Roth, R. A. (1999). Regulation of GLUT1 gene transcription by the serine/threonine kinase Akt1. J Biol Chem, 274(29), 20281–20286.

Benavente, C. A., & Jacobson, E. L. (2008). Niacin restriction upregulates NADPH oxidase and reactive oxygen species (ROS) in human keratinocytes. Free Radic Biol Med, 44(4), 527–537. doi: 10.1016/j.freeradbiomed.2007.10.006

Berger, F., Lau, C., Dahlmann, M., & Ziegler, M. (2005). Subcellular compartmentation and differential catalytic properties of the three human nicotinamide mononucleotide adenylyltransferase isoforms. J Biol Chem, 280(43), 36334–36341. doi: 10.1074/jbc.M508660200

Bi, T. Q., & Che, X. M. (2010). Nampt/PBEF/visfatin and cancer. Cancer Biol Ther, 10(2), 119–125. doi: 10.4161/cbt.10.2.12581

Cairns, R. A., Harris, I., McCracken, S., & Mak, T. W. (2011). Cancer cell metabolism. Cold Spring Harb Symp Quant Biol, 76, 299–311. doi: 10.1101/sqb.2011.76.012856

Cancer Genome Atlas Research, Network. (2008). Comprehensive genomic characterization defines human glioblastoma genes and core pathways. Nature, 455(7216), 1061–1068. doi: 10.1038/nature07385

Chiarugi, A., Dolle, C., Felici, R., & Ziegler, M. (2012). The NAD metabolome,--a key determinant of cancer cell biology. Nat Rev Cancer, 12(11), 741–752. doi: 10.1038/nrc3340

Christensen, M. K., Erichsen, K. D., Olesen, U. H., Tjornelund, J., Fristrup, P., Thougaard, A., … Bjorkling, F. (2013). Nicotinamide Phosphoribosyltransferase Inhibitors, Design, Preparation, and Structure-Activity Relationship. J Med Chem. doi: 10.1021/jm4009949

Conforti, L., Janeckova, L., Wagner, D., Mazzola, F., Cialabrini, L., Di Stefano, M., … Coleman, M. (2011). Reducing expression of NAD+ synthesizing enzyme NMNAT1 does not affect the rate of Wallerian degeneration. FEBS J, 278(15), 2666–2679. doi: 10.1111/j.1742-4658.2011.08193.x

Costanzo, M., Baryshnikova, A., Bellay, J., Kim, Y., Spear, E. D., Sevier, C. S., … Boone, C. (2010). The genetic landscape of a cell. Science, 327(5964), 425–431. doi: 10.1126/science.1180823

Cox, C., Bignell, G., Greenman, C., Stabenau, A., Warren, W., Stephens, P., … Stratton, M. R. (2005). A survey of homozygous deletions in human cancer genomes. Proc Natl Acad Sci U S A, 102(12), 4542–4547. doi: 10.1073/pnas.0408593102

DeLuna, A., Vetsigian, K., Shoresh, N., Hegreness, M., Colon-Gonzalez, M., Chao, S., & Kishony, R. (2008). Exposing the fitness contribution of duplicated genes. Nat Genet, 40(5), 676–681. doi: 10.1038/ng.123

Deshpande, R., Asiedu, M. K., Klebig, M., Sutor, S., Kuzmin, E., Nelson, J., … Myers, C. L. (2013). A comparative genomic approach for identifying synthetic lethal interactions in human cancer. Cancer Res, 73(20), 6128–6136. doi: 10.1158/0008-5472.CAN-12-3956

Dolle, C., Skoge, R. H., Vanlinden, M. R., & Ziegler, M. (2013). NAD Biosynthesis In Humans - Enzymes, Metabolites And Therapeutic Aspects. Curr Top Med Chem.

Duncan, C. G., Killela, P. J., Payne, C. A., Lampson, B., Chen, W. C., Liu, J., … Yan, H. (2010). Integrated genomic analyses identify ERRFI1 and TACC3 as glioblastoma-targeted genes. Oncotarget, 1(4), 265–277.

Forbes, S. A., Bhamra, G., Bamford, S., Dawson, E., Kok, C., Clements, J., … Stratton, M. R. (2008). The Catalogue of Somatic Mutations in Cancer (COSMIC). Curr Protoc Hum Genet, Chapter 10, Unit 10 11. doi: 10.1002/0471142905.hg1011s57

Galluzzi, L., Kepp, O., Heiden, M. G., & Kroemer, G. (2013). Metabolic targets for cancer therapy. Nat Rev Drug Discov, 12(11), 829–846. doi: 10.1038/nrd4145

Genovese, G., Ergun, A., Shukla, S. A., Campos, B., Hanna, J., Ghosh, P., … Chin, L. (2012). microRNA regulatory network inference identifies miR-34a as a novel regulator of TGF-beta signaling in glioblastoma. Cancer Discov, 2(8), 736–749. doi: 10.1158/2159-8290.CD-12-0111

Gilley, J., Adalbert, R., Yu, G., & Coleman, M. P. (2013). Rescue of peripheral and CNS axon defects in mice lacking NMNAT2. J Neurosci, 33(33), 13410–13424. doi: 10.1523/JNEUROSCI.1534-13.2013

Girolamo, M. D., Fabrizio, G., Scarpa, E. S., & Paola, S. D. (2013). NAD+-Dependent Enzymes At The Endoplasmic Reticulum. Curr Top Med Chem.

Henrich, K. O., Schwab, M., & Westermann, F. (2012). 1p36 tumor suppression--a matter of dosage? Cancer Res, 72(23), 6079–6088. doi: 10.1158/0008-5472.CAN-12-2230

Hicks, A. N., Lorenzetti, D., Gilley, J., Lu, B., Andersson, K. E., Miligan, C., … Bishop, C. E. (2012). Nicotinamide mononucleotide adenylyltransferase 2 (Nmnat2) regulates axon integrity in the mouse embryo. PLoS One, 7(10), e47869. doi: 10.1371/journal.pone.0047869

Hulsebos, T. J., Troost, D., & Leenstra, S. (2004). Molecular-genetic characterisation of gliomas that recur as same grade or higher grade tumours. J Neurol Neurosurg Psychiatry, 75(5), 723–726.

Ichimura, K., Vogazianou, A. P., Liu, L., Pearson, D. M., Backlund, L. M., Plant, K., … Collins, V. P. (2008). 1p36 is a preferential target of chromosome 1 deletions in astrocytic tumours and homozygously deleted in a subset of glioblastomas. Oncogene, 27(14), 2097–2108. doi: 10.1038/sj.onc.1210848

Jacobson, E. L., Dame, A. J., Pyrek, J. S., & Jacobson, M. K. (1995). Evaluating the role of niacin in human carcinogenesis. Biochimie, 77(5), 394–398.

Kim, D. U., Hayles, J., Kim, D., Wood, V., Park, H. O., Won, M., … Hoe, K. L. (2010). Analysis of a genome-wide set of gene deletions in the fission yeast Schizosaccharomyces pombe. Nat Biotechnol, 28(6), 617–623. doi: 10.1038/nbt.1628

Koppenol, W. H., Bounds, P. L., & Dang, C. V. (2011). Otto Warburg’s contributions to current concepts of cancer metabolism. Nat Rev Cancer, 11(5), 325–337. doi: 10.1038/nrc3038

Kotliarov, Y., Steed, M. E., Christopher, N., Walling, J., Su, Q., Center, A., … Fine, H. A. (2006). High-resolution global genomic survey of 178 gliomas reveals novel regions of copy number alteration and allelic imbalances. Cancer Res, 66(19), 9428–9436. doi: 10.1158/0008-5472.CAN-06-1691

Lal, S., Lacroix, M., Tofilon, P., Fuller, G. N., Sawaya, R., & Lang, F. F. (2000). An implantable guide-screw system for brain tumor studies in small animals. J Neurosurg, 92(2), 326–333. doi: 10.3171/jns.2000.92.2.0326

Langmead, B., & Salzberg, S. L. (2012). Fast gapped-read alignment with Bowtie 2. Nat Methods, 9(4), 357–359. doi: 10.1038/nmeth.1923

Lau, C., Niere, M., & Ziegler, M. (2009). The NMN/NaMN adenylyltransferase (NMNAT) protein family. Front Biosci (Landmark Ed), 14, 410–431.

Levine, A. J., & Puzio-Kuter, A. M. (2010). The control of the metabolic switch in cancers by oncogenes and tumor suppressor genes. Science, 330(6009), 1340–1344. doi: 10.1126/science.1193494

Martinez, R., Rohde, V., & Schackert, G. (2010). Different molecular patterns in glioblastoma multiforme subtypes upon recurrence. J Neurooncol, 96(3), 321–329. doi: 10.1007/s11060-009-9967-4

Mueller, W., Nutt, C. L., Ehrich, M., Riemenschneider, M. J., von Deimling, A., van den Boom, D., & Louis, D. N. (2007). Downregulation of RUNX3 and TES by hypermethylation in glioblastoma. Oncogene, 26(4), 583–593. doi: 10.1038/sj.onc.1209805

Muller, F. L., Colla, S., Aquilanti, E., Manzo, V. E., Genovese, G., Lee, J., … DePinho, R. A. (2012). Passenger deletions generate therapeutic vulnerabilities in cancer. Nature, 488(7411), 337–342. doi: 10.1038/nature11331

Munirajan, A. K., Ando, K., Mukai, A., Takahashi, M., Suenaga, Y., Ohira, M., … Nakagawara, A. (2008). KIF1Bbeta functions as a haploinsufficient tumor suppressor gene mapped to chromosome 1p36.2 by inducing apoptotic cell death. J Biol Chem, 283(36), 24426–24434. doi: 10.1074/jbc.M802316200

Nikiforov, A., Dolle, C., Niere, M., & Ziegler, M. (2011). Pathways and subcellular compartmentation of NAD biosynthesis in human cells: from entry of extracellular precursors to mitochondrial NAD generation. J Biol Chem, 286(24), 21767–21778. doi: 10.1074/jbc.M110.213298

Ohira, M., Kageyama, H., Mihara, M., Furuta, S., Machida, T., Shishikura, T., … Nakagawara, A. (2000). Identification and characterization of a 500-kb homozygously deleted region at 1p36.2-p36.3 in a neuroblastoma cell line. Oncogene, 19(37), 4302–4307. doi: 10.1038/sj.onc.1203786

Orsomando, G., Cialabrini, L., Amici, A., Mazzola, F., Ruggieri, S., Conforti, L., … Magni, G. (2012). Simultaneous single-sample determination of NMNAT isozyme activities in mouse tissues. PLoS One, 7(12), e53271. doi: 10.1371/journal.pone.0053271

Paik, J. H., Ding, Z., Narurkar, R., Ramkissoon, S., Muller, F., Kamoun, W. S., … DePinho, R. A. (2009). FoxOs cooperatively regulate diverse pathways governing neural stem cell homeostasis. Cell Stem Cell, 5(5), 540–553. doi: 10.1016/j.stem.2009.09.013

Petrelli, R., Felczak, K., & Cappellacci, L. (2011). NMN/NaMN adenylyltransferase (NMNAT) and NAD kinase (NADK) inhibitors: chemistry and potential therapeutic applications. Curr Med Chem, 18(13), 1973–1992.

Sahin, E., Colla, S., Liesa, M., Moslehi, J., Muller, F. L., Guo, M., … DePinho, R. A. (2011). Telomere dysfunction induces metabolic and mitochondrial compromise. Nature, 470(7334), 359–365. doi: 10.1038/nature09787

Schlisio, S., Kenchappa, R. S., Vredeveld, L. C., George, R. E., Stewart, R., Greulich, H., … Kaelin, W. G., Jr. (2008). The kinesin KIF1Bbeta acts downstream from EglN3 to induce apoptosis and is a potential 1p36 tumor suppressor. Genes Dev, 22(7), 884–893. doi: 10.1101/gad.1648608

Schulze, A., & Harris, A. L. (2012). How cancer metabolism is tuned for proliferation and vulnerable to disruption. Nature, 491(7424), 364–373. doi: 10.1038/nature11706

Shames, D. S., Elkins, K., Walter, K., Holcomb, T., Du, P., Mohl, D., … Belmont, L. D. (2013). Loss of NAPRT1 Expression by Tumor-Specific Promoter Methylation Provides a Novel Predictive Biomarker for NAMPT Inhibitors. Clin Cancer Res. doi: 10.1158/1078-0432.CCR-13-1186

Sheehan, K. M., Calvert, V. S., Kay, E. W., Lu, Y., Fishman, D., Espina, V., … Wulfkuhle, J. D. (2005). Use of reverse phase protein microarrays and reference standard development for molecular network analysis of metastatic ovarian carcinoma. Mol Cell Proteomics, 4(4), 346–355. doi: 10.1074/mcp.T500003-MCP200

Stefano, G. D., Manerba, M., & Vettraino, M. (2013). NAD Metabolism And Functions: A Common Therapeutic Target For Neoplastic, Metabolic And Neurodegenerative Diseases. Curr Top Med Chem.

Tedeschi, P. M., Markert, E. K., Gounder, M., Lin, H., Dvorzhinski, D., Dolfi, S. C., … Vazquez, A. (2013). Contribution of serine, folate and glycine metabolism to the ATP, NADPH and purine requirements of cancer cells. Cell Death Dis, 4, e877. doi: 10.1038/cddis.2013.393

Thakur, B. K., Dittrich, T., Chandra, P., Becker, A., Lippka, Y., Selvakumar, D., … Welte, K. (2012). Inhibition of NAMPT pathway by FK866 activates the function of p53 in HEK293T cells. Biochem Biophys Res Commun, 424(3), 371–377. doi: 10.1016/j.bbrc.2012.06.075

Trapnell, C., Roberts, A., Goff, L., Pertea, G., Kim, D., Kelley, D. R., … Pachter, L. (2012). Differential gene and transcript expression analysis of RNA-seq experiments with TopHat and Cufflinks. Nat Protoc, 7(3), 562–578. doi: 10.1038/nprot.2012.016

Vettraino, M., Manerba, M., Govoni, M., & Di Stefano, G. (2013). Galloflavin suppresses lactate dehydrogenase activity and causes MYC downregulation in Burkitt lymphoma cells through NAD/NADH-dependent inhibition of sirtuin-1. Anticancer Drugs, 24(8), 862–870. doi: 10.1097/CAD.0b013e328363ae50

Wiedemeyer, R., Brennan, C., Heffernan, T. P., Xiao, Y., Mahoney, J., Protopopov, A., … Chin, L. (2008). Feedback circuit among INK4 tumor suppressors constrains human glioblastoma development. Cancer Cell, 13(4), 355–364. doi: 10.1016/j.ccr.2008.02.010

Wise, D. R., DeBerardinis, R. J., Mancuso, A., Sayed, N., Zhang, X. Y., Pfeiffer, H. K., … Thompson, C. B. (2008). Myc regulates a transcriptional program that stimulates mitochondrial glutaminolysis and leads to glutamine addiction. Proc Natl Acad Sci U S A, 105(48), 18782–18787. doi: 10.1073/pnas.0810199105

Xu, Z., Du, P., Meiser, P., & Jacob, C. (2012). Proanthocyanidins: oligomeric structures with unique biochemical properties and great therapeutic promise. Nat Prod Commun, 7(3), 381–388.

Yin, D., Ogawa, S., Kawamata, N., Tunici, P., Finocchiaro, G., Eoli, M., … Koeffler, H. P. (2009). High-resolution genomic copy number profiling of glioblastoma multiforme by single nucleotide polymorphism DNA microarray. Mol Cancer Res, 7(5), 665–677. doi: 10.1158/1541-7786.MCR-08-0270

Ying, H., Kimmelman, A. C., Lyssiotis, C. A., Hua, S., Chu, G. C., Fletcher-Sananikone, E., … DePinho, R. A. (2012). Oncogenic Kras maintains pancreatic tumors through regulation of anabolic glucose metabolism. Cell, 149(3), 656–670. doi: 10.1016/j.cell.2012.01.058

Ying, H., Zheng, H., Scott, K., Wiedemeyer, R., Yan, H., Lim, C., … DePinho, R. A. (2010). Mig-6 controls EGFR trafficking and suppresses gliomagenesis. Proc Natl Acad Sci U S A, 107(15), 6912–6917. doi: 10.1073/pnas.0914930107

Yuan, M., Breitkopf, S. B., Yang, X., & Asara, J. M. (2012). A positive/negative ion-switching, targeted mass spectrometry-based metabolomics platform for bodily fluids, cells, and fresh and fixed tissue. Nat Protoc, 7(5), 872–881. doi: 10.1038/nprot.2012.024

Zhai, R. G., Cao, Y., Hiesinger, P. R., Zhou, Y., Mehta, S. Q., Schulze, K. L., … Bellen, H. J. (2006). Drosophila NMNAT maintains neural integrity independent of its NAD synthesis activity. PLoS Biol, 4(12), e416. doi: 10.1371/journal.pbio.0040416

Zois, C. E., Favaro, E., & Harris, A. L. (2014). Glycogen metabolism in cancer. Biochem Pharmacol. doi: 10.1016/j.bcp.2014.09.001

