## Supplementary Figures for "NAD+ biosynthesis as a collateral lethality target for precision oncology"

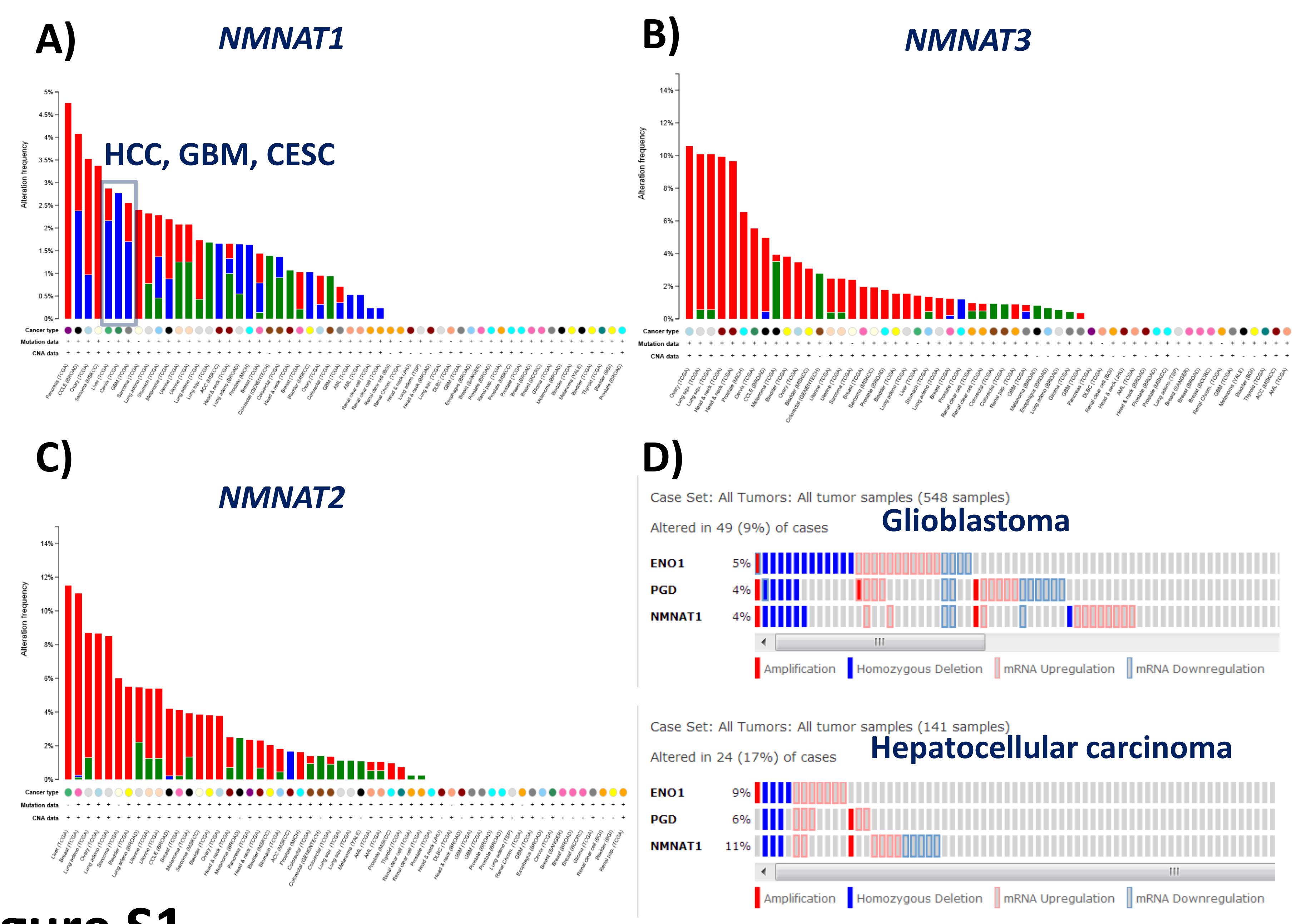

A)
NMNAT1
HCC, GBM, CESC
Alteration frequency
Cancer type
Mutation data
CNA data
B)
NMNAT3
Alteration frequency
C)
NMNAT2
Alteration frequency
D)
Case Set: All Tumors: All tumor samples (548 samples)
Altered in 49 (9%) of cases
Glioblastoma
ENO1 5%
PGD 4%
NMNAT1 4%
Amplification
Homozygous Deletion
mRNA Upregulation
mRNA Downregulation
Case Set: All Tumors: All tumor samples (141 samples)
Altered in 24 (17%) of cases
Hepatocellular carcinoma
ENO1 9%
PGD 6%
NMNAT1 11%
Amplification
Homozygous Deletion
mRNA Upregulation
mRNA Downregulation


### Figure S1

**Figure S1: Genomic alterations of NAD synthesizing genes in the Cancer Genome.** Genetic changes (point mutations, deletions, amplifications) in NAD synthesizing enzymes were queried in major human cancers using cBio portal. The genes *QPRT*, *NAPRT1*, *NAMPT* and *NADSYN1* showed non-recurrent point mutations as well as some amplifications but not genomic deletions (not shown). The situation is similar for *NMNAT2* and *3* (**C, B**), showing either amplification (Red) or point mutations (Green); only *NMNAT1*(**A**) shows consistent homozygous deletion (Blue) in diverse cancers. Deletion of NMNAT1 in Glioblastoma (GBM), Hepatocellular carcinoma (LIHC) and Cervical Squamous Carcinoma (CESC) are pointed out in a blue outline (A). Individual cases of homozygous deletion are pointed out in GBM and LIHC, showing that NMNAT1 deletion frequently occurs concurrently with ENO1 on the 1p36 locus (**D**).

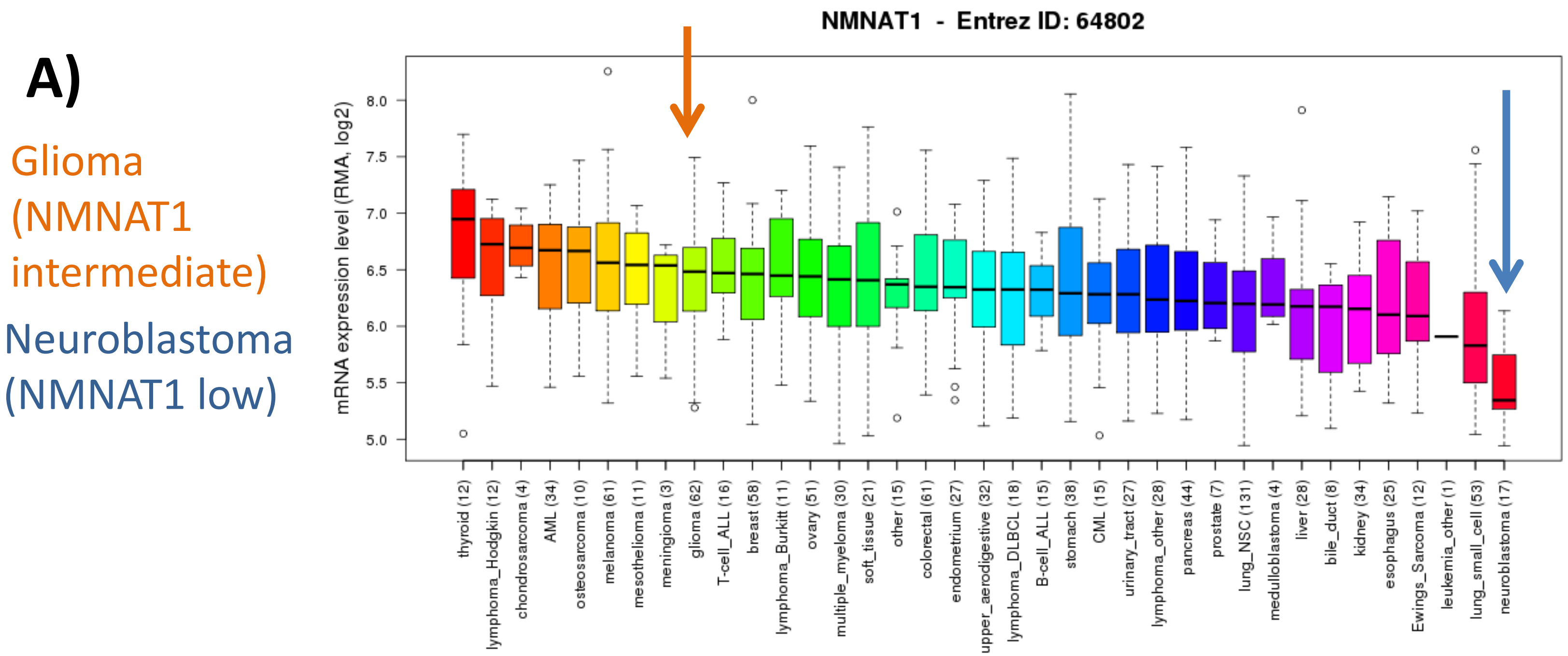
A)
Glioma
(NMNAT1
intermediate)
Neuroblastoma
(NMNAT1 low)
NMNAT1 - Entrez ID: 64802
mRNA expression level (RMA, log2)
8.0
7.5
7.0
6.5
6.0
5.5
5.0
thyroid (12)
lymphoma_Hodgkin (12)
chondrosarcoma (4)
AML (34)
osteosarcoma (10)
melanoma (61)
mesothelioma (11)
meningioma (3)
glioma (62)
T-cell_ALL (16)
breast (58)
lymphoma_Burkitt (11)
ovary (51)
multiple_myeloma (30)
soft_tissue (21)
other (15)
colorectal (61)
endometrium (27)
upper_aerodigestive (32)
lymphoma_DLBCL (18)
B-cell_ALL (15)
stomach (38)
CML (15)
urinary_tract (27)
lymphoma_other (28)
pancreas (44)
prostate (7)
lung_NSC (131)
medulloblastoma (4)
liver (28)
bile_duct (8)
kidney (34)
esophagus (25)
Ewings_Sarcoma (12)
leukemia_other (1)
lung_small_cell (53)
neuroblastoma (17)

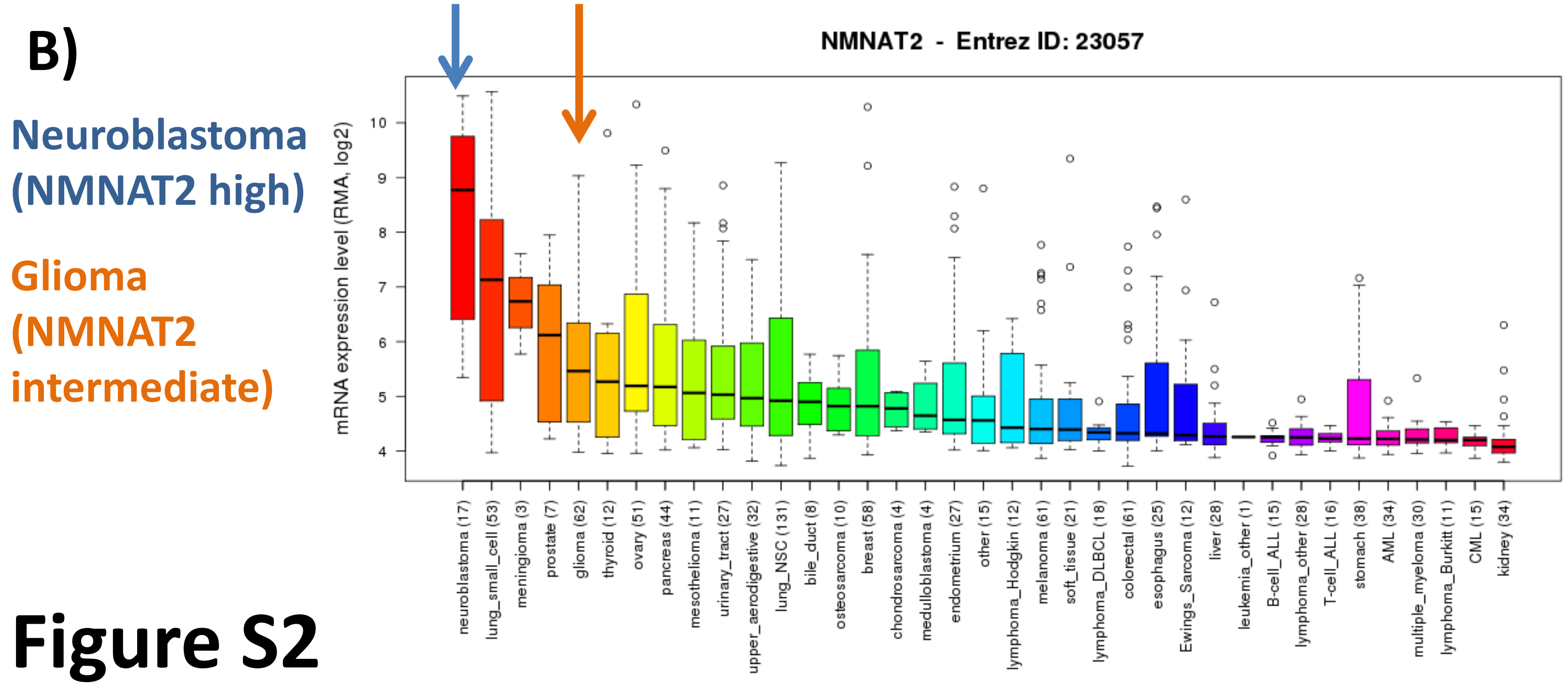
B)
Neuroblastoma
(NMNAT2 high)
Glioma
(NMNAT2
intermediate)
NMNAT2 - Entrez ID: 23057
mRNA expression level (RMA, log2)
10
9
8
7
6
5
4
neuroblastoma (17)
lung_small_cell (53)
meningioma (3)
prostate (7)
glioma (62)
thyroid (12)
ovary (51)
pancreas (44)
mesothelioma (11)
urinary_tract (27)
upper_aerodigestive (32)
lung_NSC (131)
bile_duct (8)
osteosarcoma (10)
breast (58)
chondrosarcoma (4)
medulloblastoma (4)
endometrium (27)
other (15)
lymphoma_Hodgkin (12)
melanoma (61)
soft_tissue (21)
lymphoma_DLBCL (18)
colorectal (61)
esophagus (25)
Ewings_Sarcoma (12)
liver (28)
leukemia_other (1)
B-cell_ALL (15)
lymphoma_other (28)
T-cell_ALL (16)
stomach (38)
AML (34)
multiple_myeloma (30)
lymphoma_Burkitt (11)
CML (15)
kidney (34)

### Figure S2

**Figure S2: High expression of NMNAT2 and low expression of NMNAT1 in cancer cell lines of neural origin.** Affymetrix microarray expression data for NMNAT1 and NMNAT2 was retrieved from the Cancer Cell Line Encyclopedia (Barretina et al., 2012, Nature). Expression of NMNAT1 (A) and NMNAT2 (B) were grouped and averaged in cell lines from the same tissue of origin and are shown as bar graphs in descending order. Consistent with its strong neural expression, cell lines of neuroectodermal origin exhibit high levels of NMNAT2, with the highest expression seen in neuroblastoma cell lines (B, blue arrows). Conversely, neuroblastoma cell lines also show the lowest expression of NMNAT1 (A, blue arrows). Glioma cell lines show high to intermediate expression of NMNAT1 and 2 (A, B orange arrows). The expression of NMNAT3 is low across cell lines, except those of hematopoietic origin (data not shown).

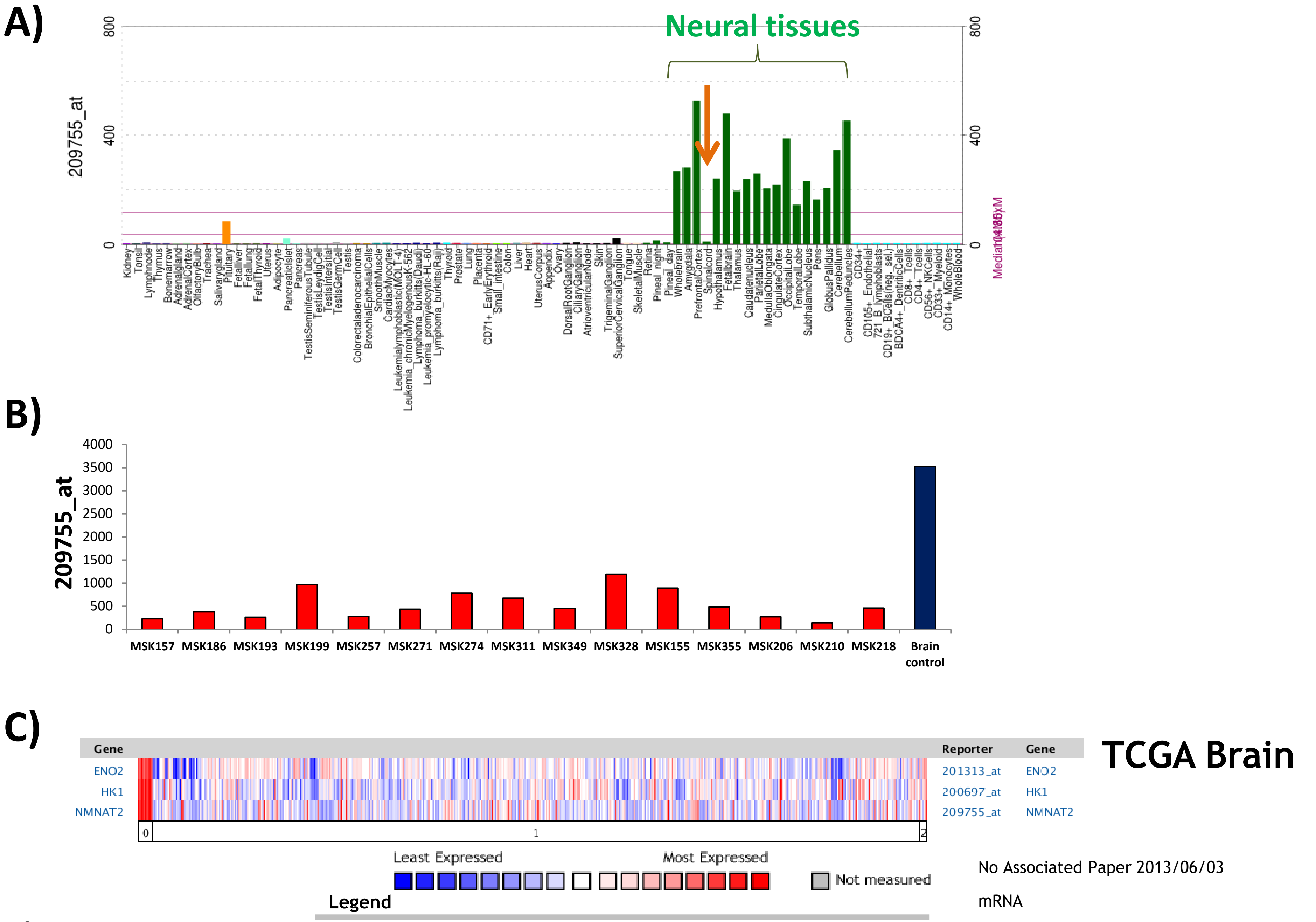
A)
Neural tissues
209755_at
800
400
0
B)
209755_at
4000
3500
3000
2500
2000
1500
1000
500
0
MSK157
MSK186
MSK193
MSK199
MSK257
MSK271
MSK274
MSK311
MSK349
MSK328
MSK155
MSK355
MSK206
MSK210
MSK218
Brain control
C)
Gene
ENO2
HK1
NMNAT2
Reporter
201313_at
200697_at
209755_at
TCGA Brain
Least Expressed
Most Expressed
Not measured
Legend
0. Normal brain (10) 1. Brain Glioblastoma (542) 2. Glioblastoma (5)
No Associated Paper 2013/06/03
mRNA
Human Genome U133A Array

### Figure S3

**Figure S3: Low expression of NMNAT2 in GBM tumors as compared to normal brain.** AffymetrixHG-U133 microarray expression data for NMNAT2 in normal human tissue was retrieved from BioGPS.org (A). Expression of NMNAT2 was highest in neural tissues, or neural derived tissues. Within neural tissues, those with the highest levels of white matter (e.g. spinal cord) had the lowest level of NMNAT2 expression, suggesting low expression in glial cells and high expression in neuronal cell bodies. Consistent with the glial origin of GBM, querying our in-house generated Affymetrix microarray data (B, GSE9200, (Wiedemeyer et al., 2008, Cancer Cell)) reveals that primary GBM tumors have low expression of NMNAT2 compared to normal brain, a conclusion supported by other public domain microarray studies, such as U133A microarray data for TCGA GBM, C (The Oncomine™ Platform (Life Technologies, Ann Arbor, MI).

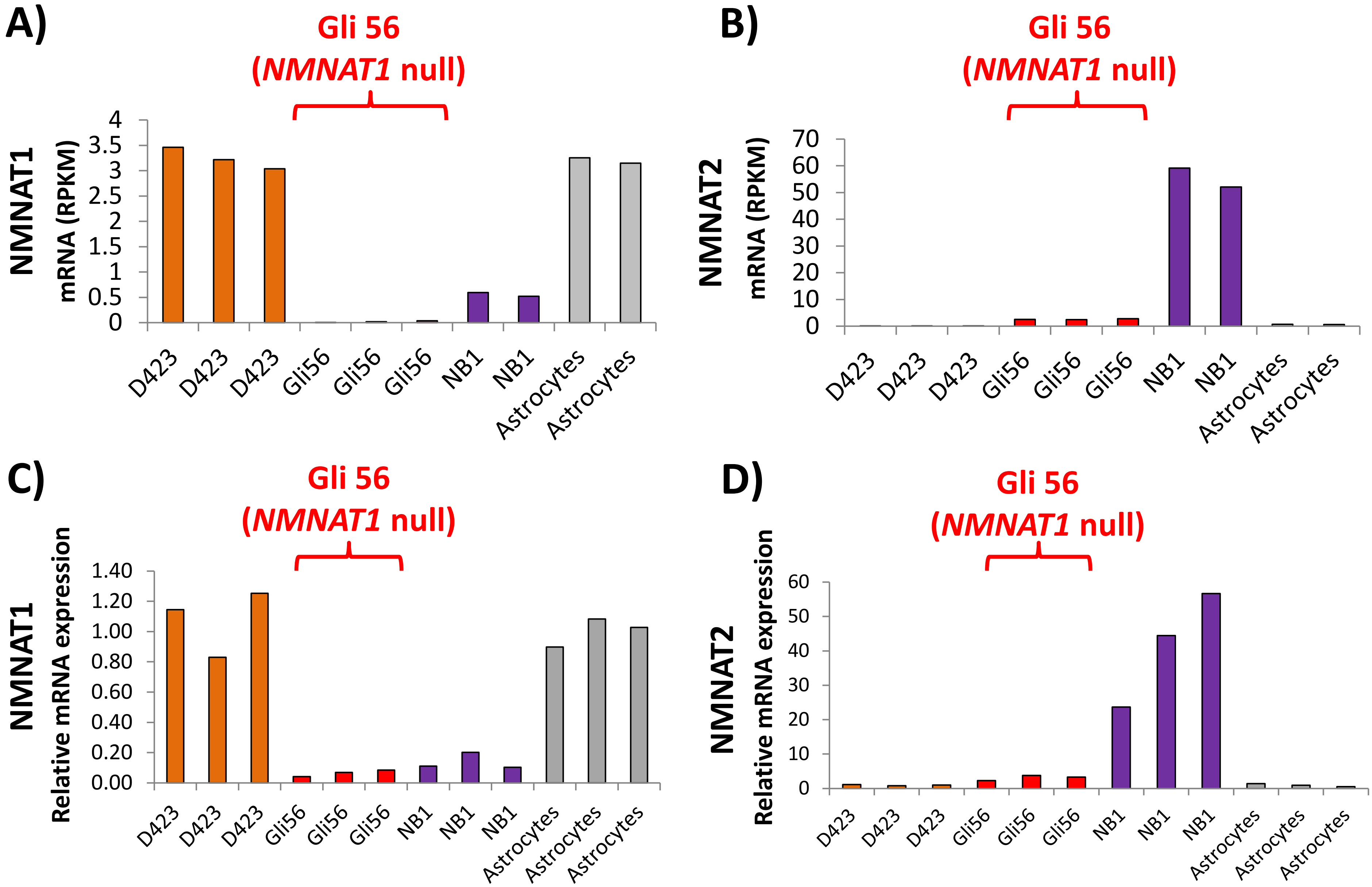
A)
Gli 56
(NMNAT1 null)
NMNAT1
mRNA (RPKM)
4
3.5
3
2.5
2
1.5
1
0.5
0
D423
D423
D423
Gli56
Gli56
Gli56
NB1
NB1
Astrocytes
Astrocytes
B)
Gli 56
(NMNAT1 null)
NMNAT2
mRNA (RPKM)
70
60
50
40
30
20
10
0
D423
D423
D423
Gli56
Gli56
Gli56
NB1
NB1
Astrocytes
Astrocytes
C)
Gli 56
(NMNAT1 null)
NMNAT1
Relative mRNA expression
1.40
1.20
1.00
0.80
0.60
0.40
0.20
0.00
D423
D423
D423
Gli56
Gli56
Gli56
NB1
NB1
NB1
Astrocytes
Astrocytes
Astrocytes
D)
Gli 56
(NMNAT1 null)
NMNAT2
Relative mRNA expression
60
50
40
30
20
10
0
D423
D423
D423
Gli56
Gli56
Gli56
NB1
NB1
NB1
Astrocytes
Astrocytes
Astrocytes

### Figure S4

**Figure S4: NMNAT1 and NMNAT2 expression in normal astrocytes and 1p36 homozygously deleted cancer cell lines.** NMNAT1 (A, C) and NMNAT2 (B, D) expression was determined by RNA-seq (A, B) and Q-PCR (C, D) in the D423-MG (ENO1) and Gli56 (ENO1 NMANT1 null) glioma cell lines as well as the NB1 neuroblastoma cell line and normal human astrocytes. In line with the known pattern of high expression of NMNAT2 expression in neural tissue, NB1 neuroblastoma cells showed the highest levels of the NMNAT2 expression. Gli56 *NMNAT1*-null cells showed ~2.5-fold higher level of expression compared to astrocytes and D423 glioma cells, suggesting some degree of compensatory upregulation, but this fell well below the levels seen in NB1 cells.

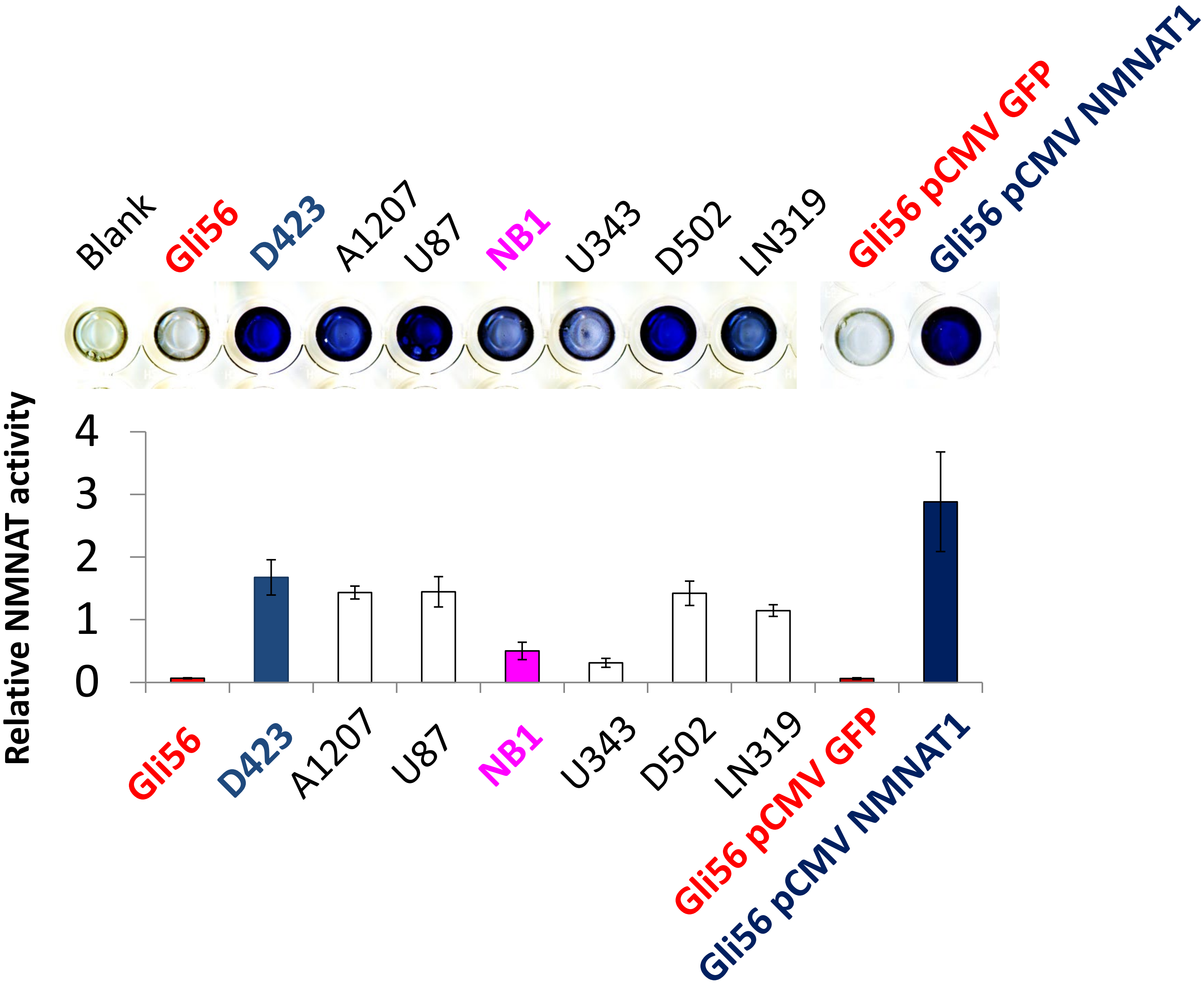
Blank
Gli56
D423
A1207
U87
NB1
U343
D502
LN319
Gli56 pCMV GFP
Gli56 pCMV NMNAT1
Relative NMNAT activity
4
3
2
1
0
Gli56
D423
A1207
U87
NB1
U343
D502
LN319
Gli56 pCMV GFP
Gli56 pCMV NMNAT1

### Figure S5

**Figure S5: Exceptionally low NMNAT activity in *NMNAT1*-deleted glioma cells.** NMNAT activity was measured spectrophotometrically in native lysates in a panel of cancer cell lines, including GLI56 *NMNAT1*-deleted (Red bars), *NMNAT1*-rescued (Blue bars) and NB1 *NMNAT1*-WT cells (Pink bars). Activity was normalized across cell lines and expressed as a ratio to the average activity. Each bar represents the average of three replicates +/- S.D. Background (blank) is yellow (MTT), with high NMNAT activity samples turning dark blue. Gli56 *NMNAT1*-deleted cells (red bars) showed exceptionally low NMNAT activity, which was completely restored by ectopic re-expression of *NMNAT1* (blue bars).

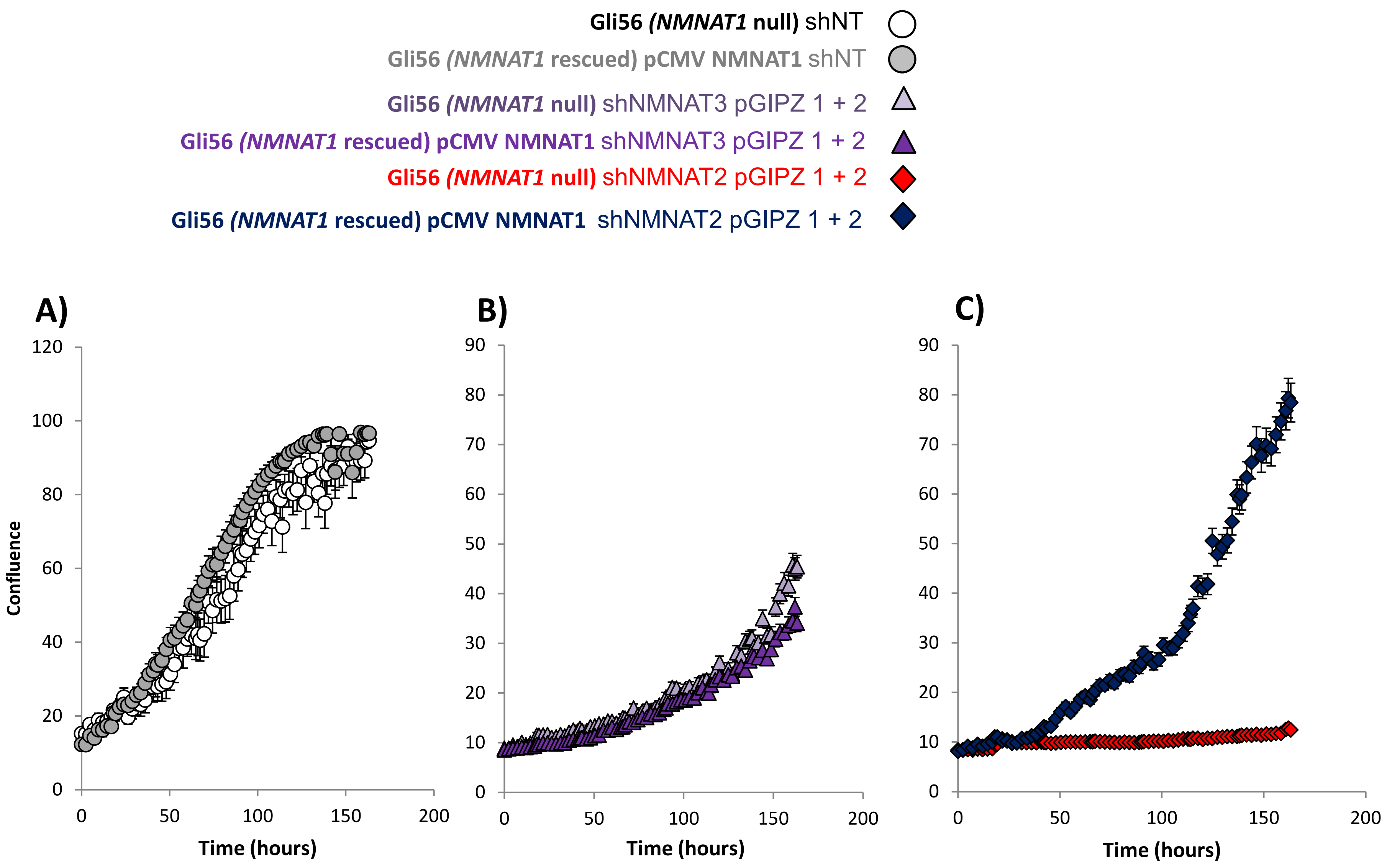

Gli56 (NMNAT1 null) shNT
Gli56 (NMNAT1 rescued) pCMV NMNAT1 shNT
Gli56 (NMNAT1 null) shNMNAT3 pGIPZ 1 + 2
Gli56 (NMNAT1 rescued) pCMV NMNAT1 shNMNAT3 pGIPZ 1 + 2
Gli56 (NMNAT1 null) shNMNAT2 pGIPZ 1 + 2
Gli56 (NMNAT1 rescued) pCMV NMNAT1 shNMNAT2 pGIPZ 1 + 2
A)
B)
C)
Confluence
Time (hours)
Time (hours)
Time (hours)


### Figure S6

**Figure S6: Stable shRNA knockdown of NMNAT2 selectively stalls the growth of *NMNAT1* null but not *NMNAT1* rescued glioma cells.**
Parental Gli56 *NMNAT1* null cells or Gli56 rescued by ectopic re-expression were infected with pGIPZ non-targeting (A), shNMNAT3 1 + 2 (B), and shNMNAT2 1 + 2 (C), and growth was followed by live-imaging using the Incucyte. Each trace shows the average of four wells, +/- S.E.M. ShRNA mediated ablation of NMNAT2 dramatically inhibited the growth of Gli56 NMNAT1 null, but not NMNAT1 rescued cells. NMNAT3 knockdown inhibited growth of both cell lines.

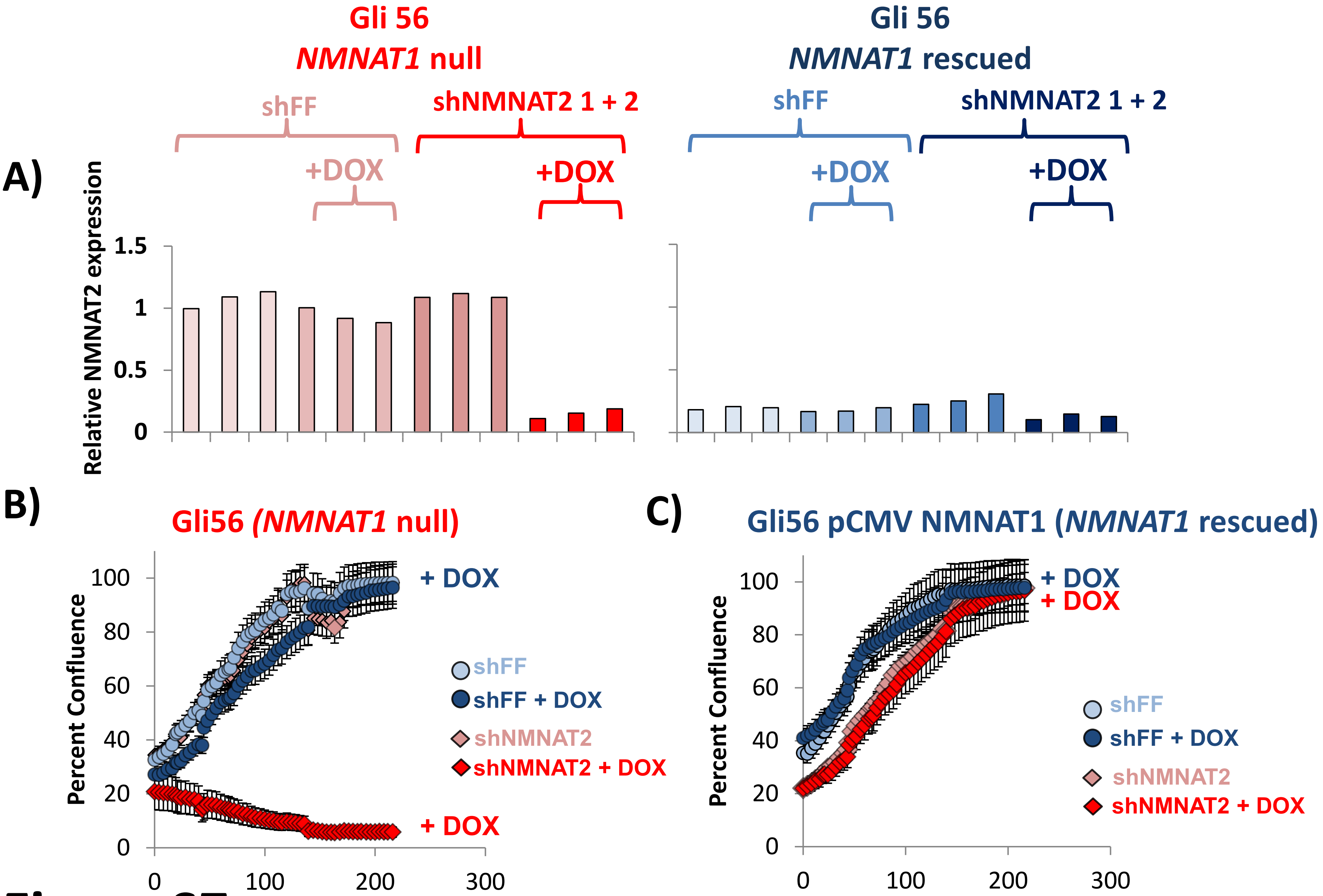
A)
Gli 56
NMNAT1 null
shFF
shNMNAT2 1 + 2
+DOX
+DOX
Gli 56
NMNAT1 rescued
shFF
shNMNAT2 1 + 2
+DOX
+DOX
Relative NMNAT2 expression
1.5
1
0.5
0
B)
Gli56 (NMNAT1 null)
Percent Confluence
100
80
60
40
20
0
0
100
200
300
Time (hours)
+ DOX
+ DOX
shFF
shFF + DOX
shNMNAT2
shNMNAT2 + DOX
C)
Gli56 pCMV NMNAT1 (NMNAT1 rescued)
Percent Confluence
100
80
60
40
20
0
0
100
200
300
Time (hours)
+ DOX
+ DOX
shFF
shFF + DOX
shNMNAT2
shNMNAT2 + DOX

### Figure S7

**Figure S7: Inducible pTRIPZ shRNA knockdown of NMNAT2 is selectively toxic to *NMNAT1*-deleted glioma cells.** Gli56 parental and Gli56 NMNAT1 rescued glioma cells were infected with pTRIPZ DOX-inducible shRNA against NMNAT2 (shNMNAT2-1+2) or non-targeting control (shFF). Knockdown of NMNAT2 was confirmed by Q-PCR (**A**); interestingly Gli56 NMNAT1 rescued cells exhibited very low NMNAT2 expression (blue to light blue bars) as compared to NMNAT1 null parental cells (red to light red bars), suggesting a complementary regulatory interaction between the two isoforms. Each bar represents an individual biological replicate. Induction of shNMNAT2 by DOX had minimal effects on growth of Gli56 NMNAT1-rescued cells (Incucyte, **C,** blue), but dramatically inhibited the growth and killed Gli56 parental NMNAT1 null cells (**B;** red). Each tracing shows the average of four wells +/- S.E.M.

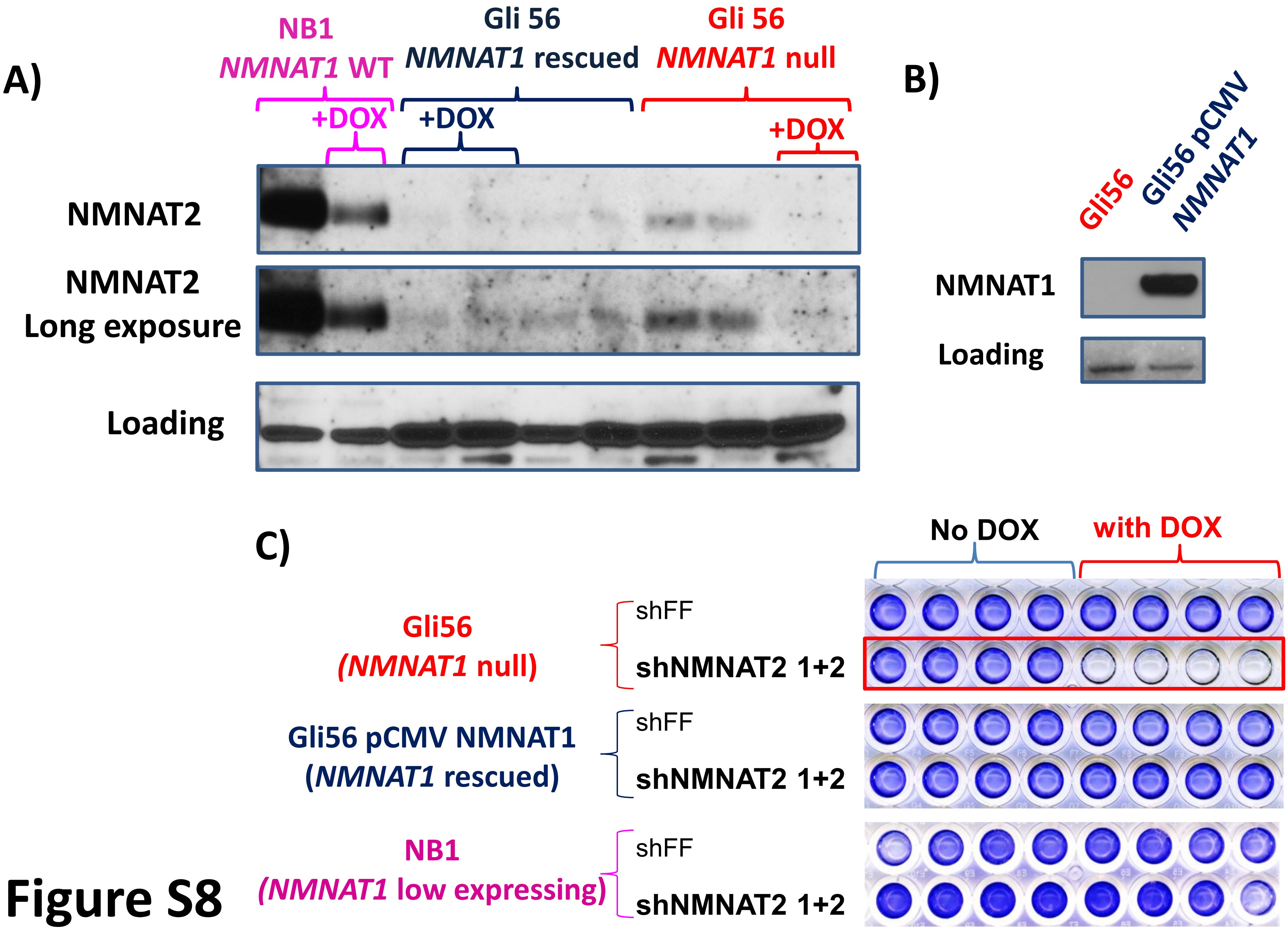
A)
NB1
NMNAT1 WT
Gli 56
NMNAT1 rescued
Gli 56
NMNAT1 null
+DOX
+DOX
+DOX
NMNAT2
NMNAT2
Long exposure
Loading
B)
Gli56
Gli56 pCMV NMNAT1
NMNAT1
Loading
C)
No DOX
with DOX
Gli56
(NMNAT1 null)
shFF
shNMNAT2 1+2
Gli56 pCMV NMNAT1
(NMNAT1 rescued)
shFF
shNMNAT2 1+2
NB1
(NMNAT1 low expressing)
shFF
shNMNAT2 1+2

**Figure S8**

**Figure S8: Potent knockdown of NMNAT2 in NB1 neuroblastoma cells has minimal phenotypic effects** Inducible shRNA (pTRIPZ shNMNAT2 1 + 2) mediated knockdown of NMNAT2 resulted in a >95% reduction in NMNAT2 protein (**A**). However, because of high expression of NMNAT2 in neuroblastoma cells, the residual NMNAT2 expression after shRNA knockdown was still higher than that found under basal conditions in the Gli56 glioblastoma cell line. Thus, despite this great knockdown, the doxycycline induction of shNMNAT2 had minimal effects on cellular growth and viability (**C**, crystal violet-stained plates).

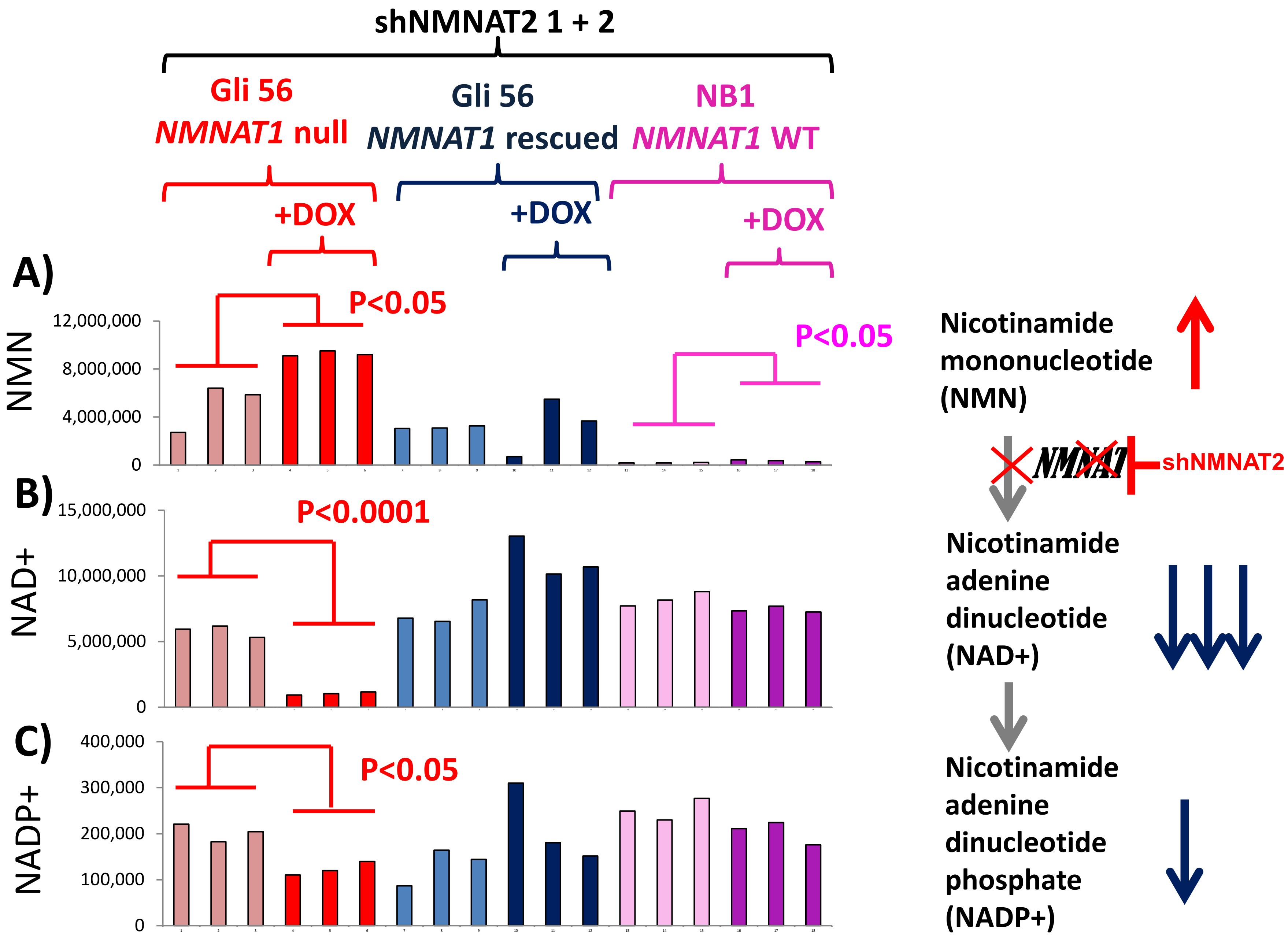
shNMNAT2 1 + 2
Gli 56
NMNAT1 null
Gli 56
NMNAT1 rescued
NB1
NMNAT1 WT
+DOX
+DOX
+DOX
A)
P<0.05
P<0.05
NMN
12,000,000
8,000,000
4,000,000
0
B)
15,000,000
10,000,000
5,000,000
0
NAD+
P<0.0001
C)
400,000
300,000
200,000
100,000
0
NADP+
P<0.05
Nicotinamide mononucleotide (NMN)
NMNAT
shNMNAT2
Nicotinamide adenine dinucleotide (NAD+)
Nicotinamide adenine dinucleotide phosphate (NADP+)

### Figure S9

**Figure S9: Inducible shRNA knockdown of NMNAT2 selectively decreases NAD+ levels.** LC-MS/MS based metabolomics via selected reaction monitoring (SRM) data were interrogated for alterations of adenine dinucleotides in response to DOX-inducible pTRIPZ shRNA NMNAT2 (shNMNAT2 1 + 2) knockdown in *NMNAT1*-deleted (Red), NMNAT1-rescued Gli56 glioma cells (Blue) and *NMNAT1*-WT NB1 cells (Purple). Each bar represents an individual biological replicate. Levels of Nicotinamide mononucleotide, the substrate of NMNAT, increased in response to NMNAT2 knockdown in both Gli56 NMNAT1 null and NB1 NMNAT1-WT cells, though in the latter case, this was from a much lower baseline (**A**). NAD+ levels were selectively decreased by NMNAT2 knockdown only in Gli56 NMNAT1 null cells (**B**). NADP+ levels followed a similar trend, but in a lower magnitude, both in absolute and relative terms (**C**).

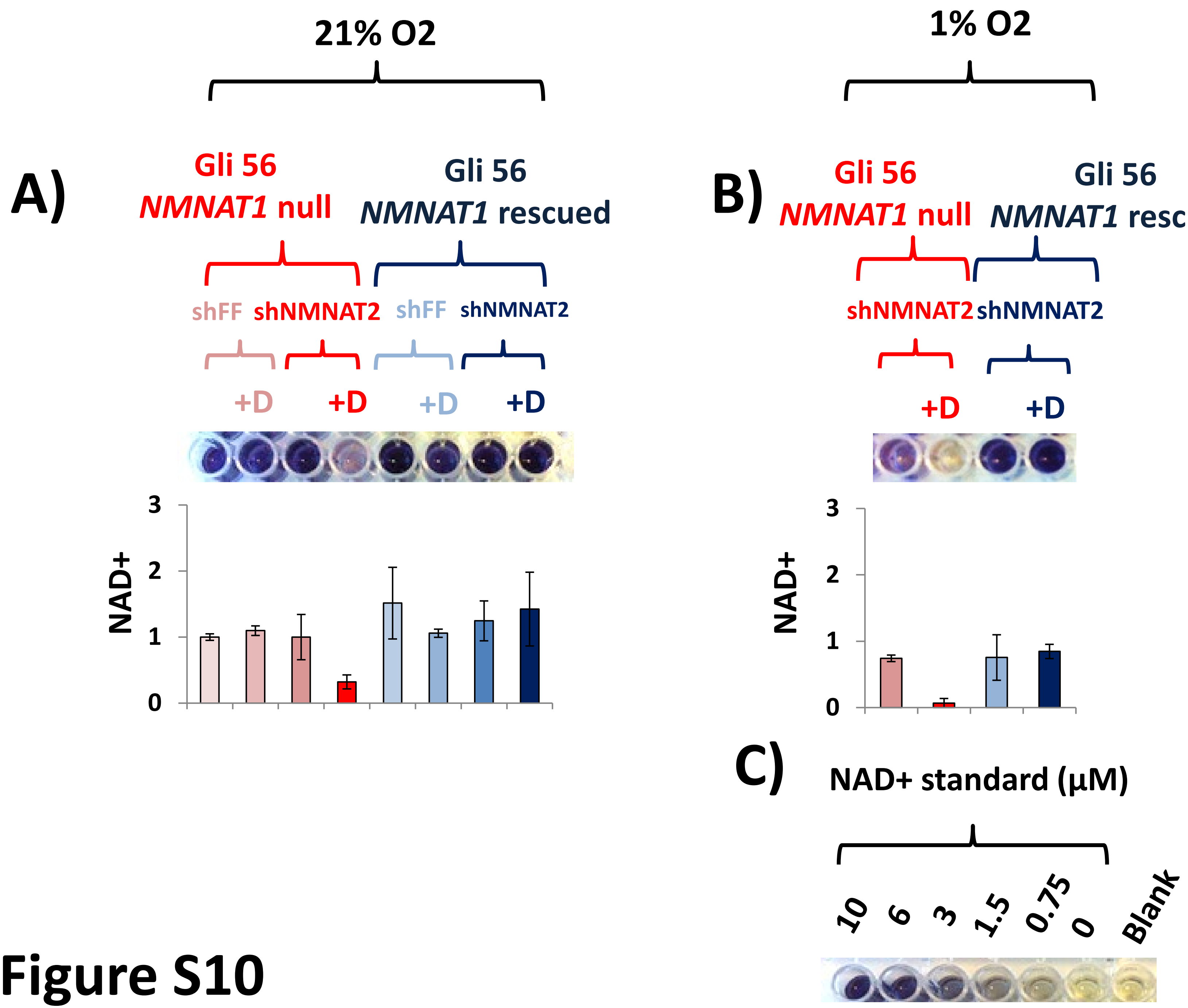
21% O2
1% O2
A)
Gli 56
NMNAT1 null
Gli 56
NMNAT1 rescued
shFF
shNMNAT2
shFF
shNMNAT2
+D
+D
+D
+D
NAD+
3
2
1
0
B)
Gli 56
NMNAT1 null
Gli 56
NMNAT1 rescued
shNMNAT2
shNMNAT2
+D
+D
NAD+
3
2
1
0
C)
NAD+ standard (µM)
10
6
3
1.5
0.75
0
Blank

### Figure S10

**Figure S10: NMNAT2 knockdown selectively decreases NAD+ levels which is exaggerated by hypoxia.** NAD+ levels were measured by an MTT-cycling assay (Enzychrom) in response to shRNA knockdown of NMNAT2 (shNMNAT2 1 + 2) in *NMNAT1*-deleted or *NMNAT1*-rescued glioma cells, under normoxic (21% $O_2$ **A**) and hypoxic conditions (1%$O_2$ **B**). Results are expressed with reference to untreated control *NMNAT1*-null glioma cells; average of three measurements +/- S.E.M. Hypoxia lowered the levels of NAD+ in all cell lines, but the effect was particularly dramatic in *NMNAT1*-null glioma cells with activated shNMNAT2.

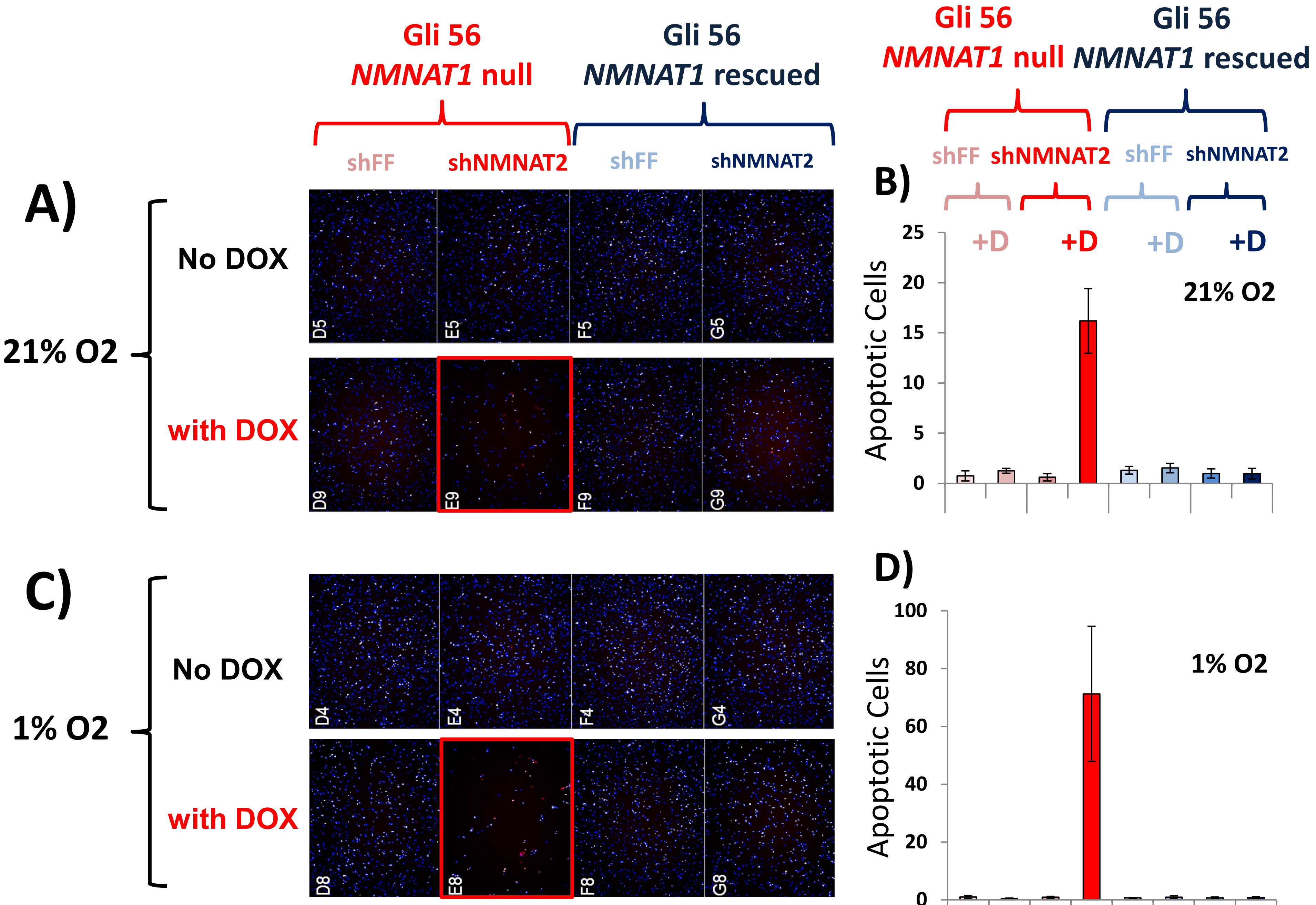
Gli 56
NMNAT1 null
Gli 56
NMNAT1 rescued
shFF
shNMNAT2
shFF
shNMNAT2
A)
21% O2
No DOX
with DOX
D5
E5
F5
G5
D9
E9
F9
G9
B)
+D
+D
+D
+D
Apoptotic Cells
21% O2
25
20
15
10
5
0
C)
1% O2
No DOX
with DOX
D4
E4
F4
G4
D8
E8
F8
G8
D)
Apoptotic Cells
1% O2
100
80
60
40
20
0

### Figure S11

**Figure S11: Selective killing by NMNAT2 knockdown is accelerated under hypoxia.** Apoptosis was followed by YO-PRO®-1 Iodide (red) and counter stained with Hoechst 33342 (blue), with the fraction of apoptotic (red) to viable cells (blue) being scored. shRNA knockdown of NMNAT2 (red, blue) or non-targeting control (shFF, light red, light blue) was induced by DOX in normoxia (21% $O_2$ **A**,**B**) and hypoxia (1% $O_2$ **C**,**D**); Representative wells are shown in **A**, **C** with averaged results of four wells +/- S.D shown in **B**,**D**. (**A**). Knockdown of NMNAT2 caused a profound induction in apoptosis specifically in *NMNAT1*-deleted (red) but not *NMNAT1*-rescued (blue) glioma cells, which was considerably exaggerated under hypoxia.

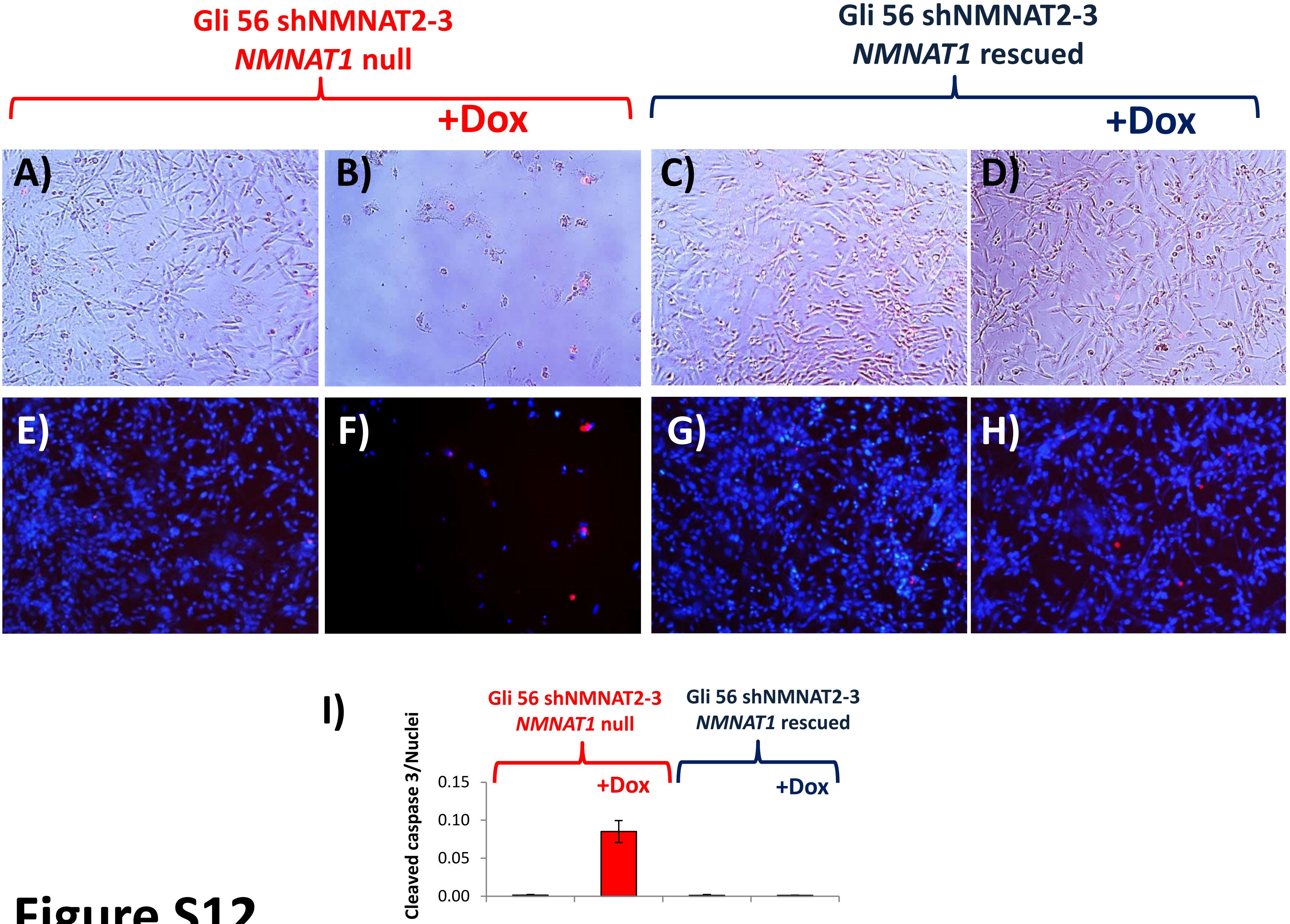
Gli 56 shNMNAT2-3
NMNAT1 null
+Dox
Gli 56 shNMNAT2-3
NMNAT1 rescued
+Dox
A)
B)
C)
D)
E)
F)
G)
H)
I)
Gli 56 shNMNAT2-3
NMNAT1 null
Gli 56 shNMNAT2-3
NMNAT1 rescued
+Dox
+Dox
Cleaved caspase 3/Nuclei
0.15
0.10
0.05
0.00

### Figure S12

**Figure S12: Knockdown of NMNAT2 leads to induction of apoptosis.**
Induction of shNMNAT2-3 by DOX in Gli56 parental*NMNAT1*-deleted glioma cells led to a potent cell killing. Immunofluorescence blotting against cleaved caspase 3 (red, nuclei stained by Hoechst 33342 are in blue) confirms induction of apoptosis (**B,F**), which was fully rescued by ectopic re-expression of NMNAT1 (**D,H**). A significant fraction of detached dead cells were positive for cleaved caspase 3, indicating death by apoptosis. Panels (**A-D**) show cleaved caspase 3 positive cells (red) and cell morphology (phase contrast). Panels (**E-F**) show total nuclei (blue, Hoechst) with cleaved caspase 3 positive cells (red) in the same fields.

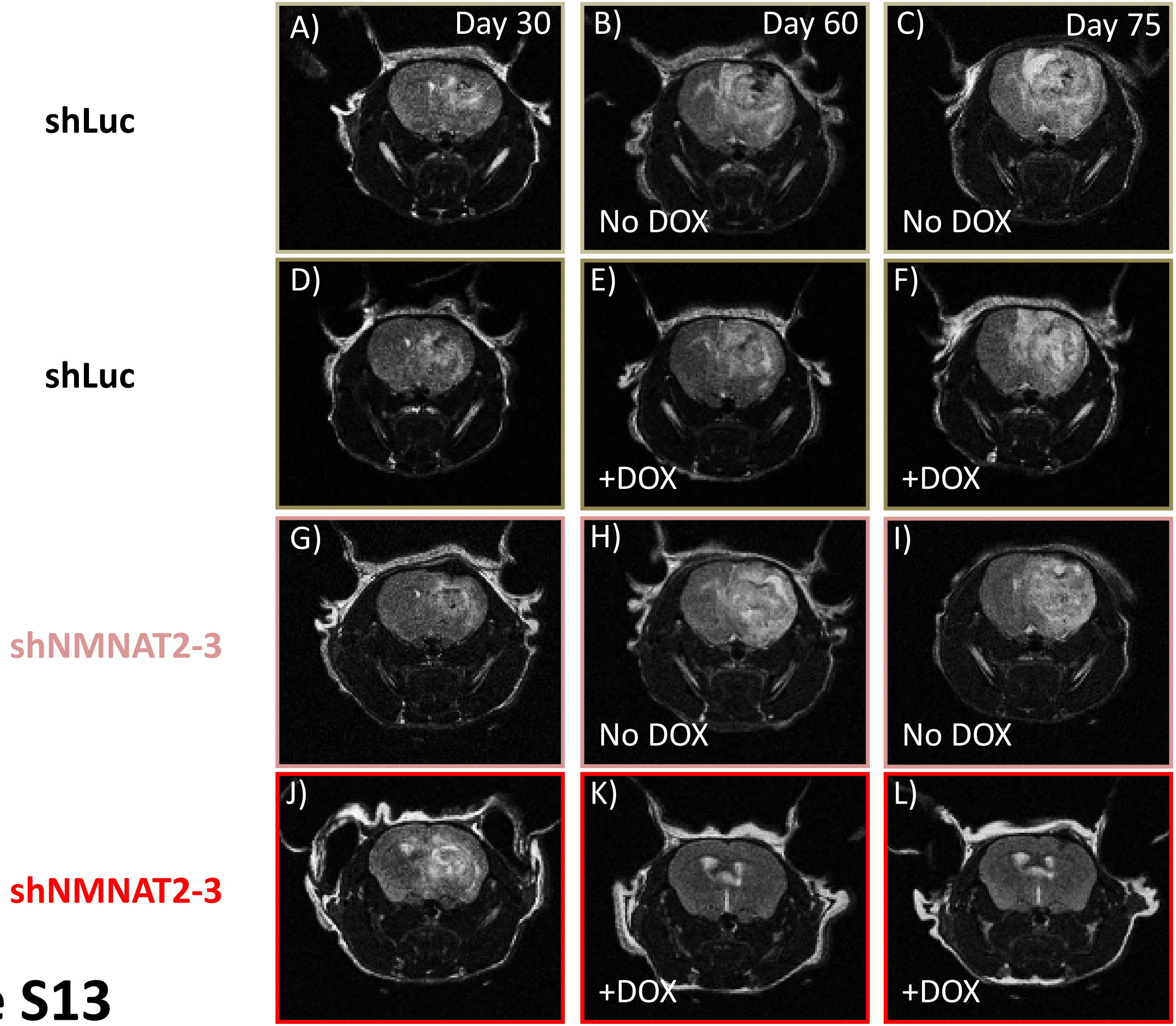
shLuc
shLuc
shNMNAT2-3
shNMNAT2-3
A)
Day 30
B)
Day 60
No DOX
C)
Day 75
No DOX
D)
E)
+DOX
F)
+DOX
G)
H)
No DOX
I)
No DOX
J)
K)
+DOX
L)
+DOX

### Figure S13

**Figure S13: Intracranial tumor eradication by induction of shRNA targeting NMNAT2 but not non-targeting control.** Gliomas were generated by intracranial injection of Gli56 NMNAT1 deleted glioma cells bearing either shRNA against NMNAT2 (G-L) or non-targeting control (A-F). Axial T2-MRI images of four individual intracranial tumor bearing mice (pictures in each row represents the same individual, at different time points), taken 30 days (A,D,G,J) after intracranial cell injection and again 30 days later (B,E,H,K) and again 15 days after that (C, F, I, L). After the initial scans, two mice (E,F and K,L) began receiving DOX to induce the shRNA. Induction of shNMNAT2 (K,L) led to a dramatic reduction in tumor area (white T2-hyperintensities) while induction of non-targeting shRNA (E,F) did not substantially alter the growth compared to non-induced (B,C).
