## Supplementary Table S1 for "NAD+ biosynthesis as a collateral lethality target for precision oncology"

| | Identity with NMNAT1 | Location | Expression | Knockout mouse phenotype | Heterozygous Knockout Phenotype | Associated Human Genetic Disease | Mean DepMap Chronos Score ~1000 cell lines (Lower numbers indicate greater lethality) |
|---|---|---|---|---|---|---|---|
| *NMNAT1* | 100% | 1p36 | Ubiquitous | Lethal <E10 | None | Leber congenital amaurosis | -0.31837 |
| *NMNAT2* | 32% | 1q25 | CNS | Post-natal lethal, but viable in combination with *Sarm1* KO | None, hypo-heterozygous with <25% residual activity are phenotypically normal | Polyneuropathy and erythromelalgia | -0.01757 |
| *NMNAT3* | 47% | 3q23 | Ubiquitous, high in blood | Viable – hemolytic anemia | None | None described | -0.04237 |

References: Loreto *et al* (2021) Neurotoxin-mediated potent activation of the axon degeneration regulator SARM1 eLife 10:e72823.

Hikosaka et al (2014) Deficiency of Nicotinamide Mononucleotide Adenylyltransferase 3 (Nmnat3) Causes Hemolytic Anemia by Altering the Glycolytic Flow in Mature Erythrocytes Journal of Biological Chemistry, Volume 289, Issue 21, 14796 - 14811

### Broad DepMap CRISPR Chronos Scores for NMNAT paralogs in a panel of ~1000 cancer cell lines shows strongest dependence on NMNAT1 with minimal dependency on NMNAT2 and 3

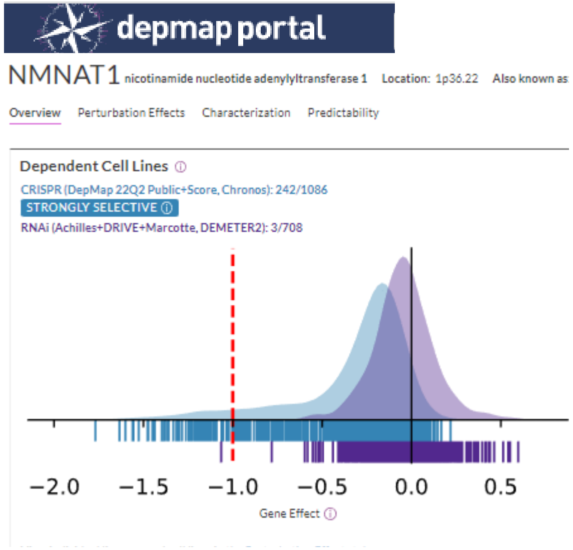


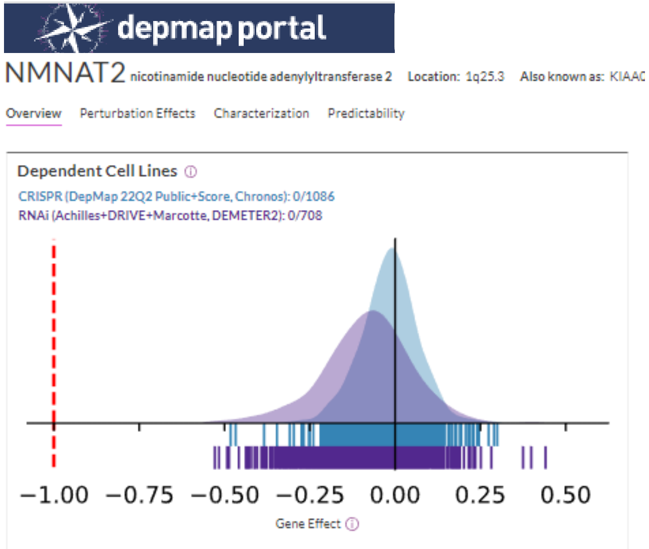


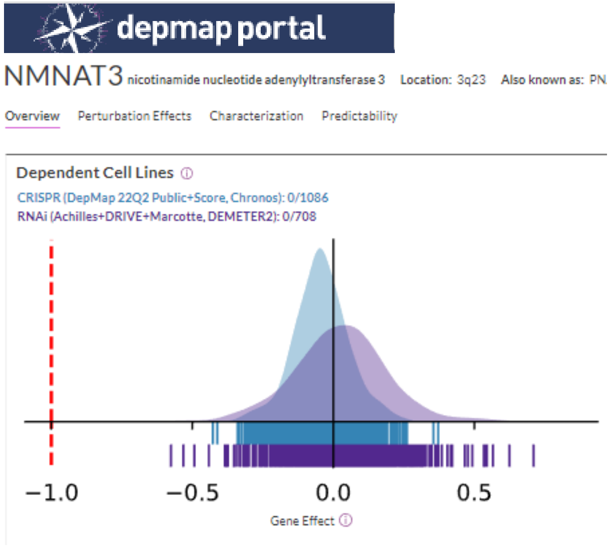


CRISPR (DepMap 22Q2 Public+Score, Chronos)

### NMNAT1 dependency is inversely correlated with expression of NMNAT2 and NMNAT3

**depmap portal**

**NMNAT1** nicotinamide nucleotide adenylyltransferase 1 Location: 1p36.22 Also known as:

Overview Perturbation Effects Characterization Predictability

Information about this page | Download data for NMNAT1 | Download data for all genes | Download input data

**CRISPR (DepMap 22Q2 Public+Score, Chronos)**

| MODEL | ACCURACY SCORE (Pearson) |
|---|---|
| Core Omics | 0.599 |

| Feature | Relative Importance | Correlation | Feature Type |
|---|---|---|---|
| IKZF1 | 10.4% | -0.55 | Expression |
| TSPAN6 | 4.1% | 0.47 | Expression |
| WAS | 3.3% | -0.53 | Expression |
| CD53 | 3.2% | -0.53 | Expression |
| MZB1 | 2.4% | -0.51 | Expression |
| RUSC2 | 2.3% | 0.33 | Expression |
| SEPTIN1 | 2.2% | -0.50 | Expression |
| FKBP9 | 2.1% | 0.46 | Expression |
| Solid | 1.9% | 0.55 | Lineage |
| TJP1 | 1.7% | 0.48 | Expression |

| MODEL | ACCURACY SCORE (Pearson) |
|---|---|
| Related | 0.619 |

| Feature | Relative Importance | Correlation | Feature Type |
|---|---|---|---|
| Solid | 24.4% | 0.55 | Lineage |
| Haematopoietic_and_Ly... | 16.6% | -0.55 | Lineage |
| Ⓡ NMNAT3 | 9.2% | 0.15 | Expression |
| cell_line_NNMD | 2.2% | -0.09 | Confounders |
| Ⓡ HMGA2 | 2.0% | 0.35 | Expression |
| Ⓡ TCEA2 | 1.5% | 0.07 | Copy num. |
| Ⓡ NMNAT2 | 1.4% | 0.27 | Expression |
| Ⓡ UBE2I | 1.3% | -0.14 | Expression |
| Ⓡ ATP6V1B2 | 1.2% | -0.11 | Expression |
| Ⓡ PARP1 | 1.1% | -0.30 | Expression |

| MODEL | ACCURACY SCORE (Pearson) |
|---|---|
| DNA-based | 0.536 |

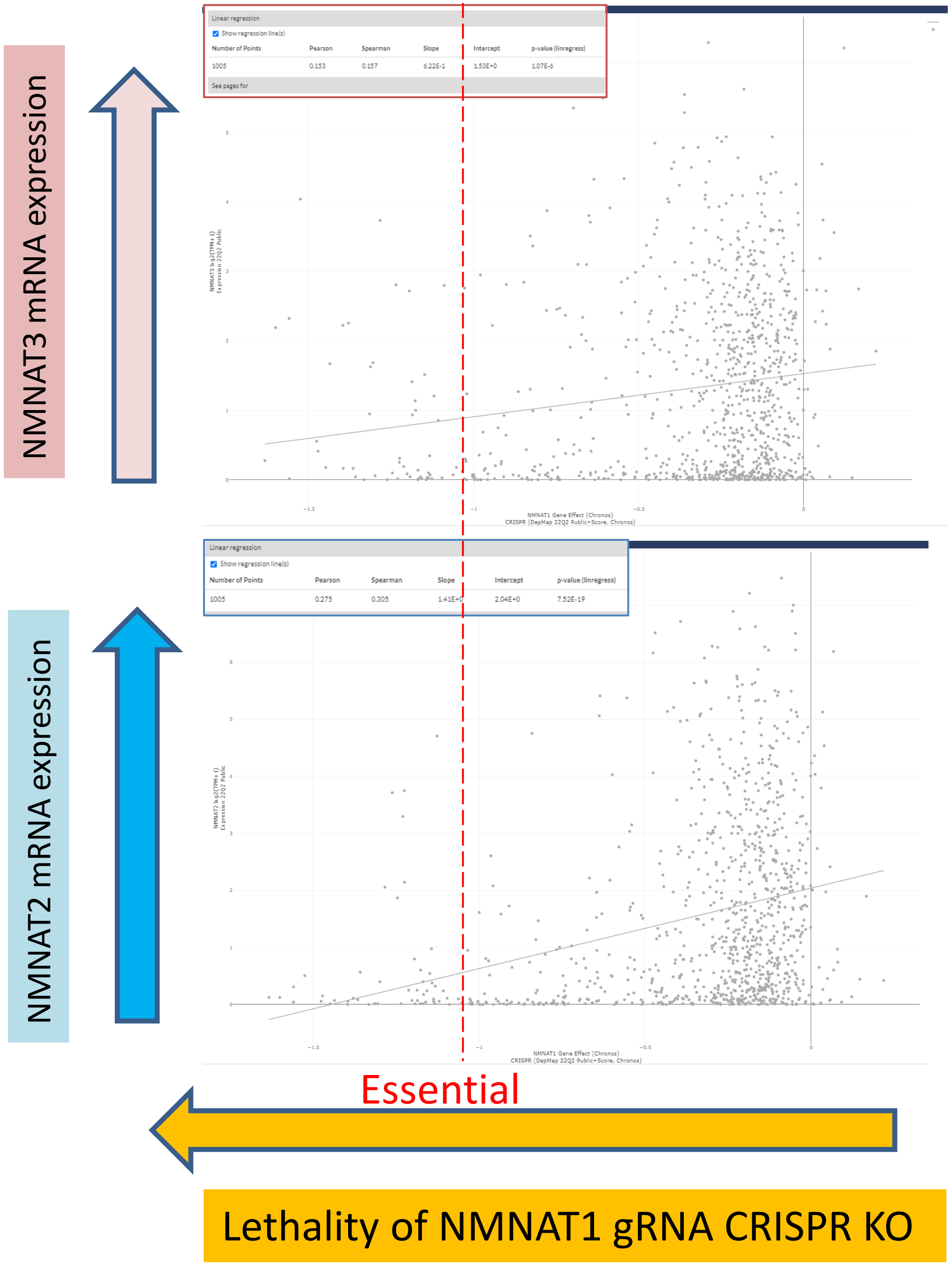

### ASCAT-corrected homozygous-deletions in COSMIC (Sanger)

| Sample | Gene | Expression | Expr Level (Z-Score) | CN Type | Minor Allele | Copy Number | CNV segment Posn. | Average Ploidy | Study | Cancer Site |
|---|---|---|---|---|---|---|---|---|---|---|
| TCGA-F4-6855-01 | NMNAT1 | Normal | -0.87 | Loss | 0 | 0 | 1:61735..121741181 | 1.98 | 376 | Colon Adenocarcinoma |
| TCGA-DD-AADJ-01 | NMNAT1 | Under | -2.80 | Loss | 0 | 0 | 1:9914107..10015782 | 3.90 | 628 | Liver Hepatocellular carcinoma |
| TCGA-W2-A7HB-01 | NMNAT1 | - | - | Loss | 0 | 0 | 1:61735..121741181 | 1.71 | 664 | Pheochromocytoma and Paraganglioma |
| TCGA-06-0879-01 | NMNAT1 | - | - | Loss | 0 | 0 | 1:7839598..10424794 | 2.03 | 329 | Glioblastoma Multiforme |
| TCGA-W5-AA2T-01 | NMNAT1 | - | - | Loss | 0 | 0 | 1:7567847..10293312 | 3.53 | 662 | Cholangiocarcinoma |
| TCGA-OR-A5LL-01 | NMNAT1 | Under | -2.24 | Loss | 0 | 1 | 1:8744829..12759130 | 3.85 | 631 | 2743554 |
| TCGA-52-7810-01 | NMNAT1 | Under | -2.07 | Loss | 0 | 1 | 1:8627807..18932229 | 3.71 | 418 | 2804522 |
| TCGA-AA-3858-01 | NMNAT1 | Under | -2.43 | Loss | 0 | 1 | 1:9460977..10203089 | 4.15 | 376 | 2048892 |
| TCGA-ZG-A9L9-01 | NMNAT1 | Under | -2.47 | Loss | 0 | 1 | 1:9438054..12696860 | 3.76 | 435 | 2463243 |
| TCGA-A8-A09X-01 | NMNAT1 | Normal | -1.52 | Loss | 0 | 1 | 1:8435885..11763589 | 3.81 | 414 | 1471981 |

### TCGA RNA data identify tumors with exceptionally low NMNAT1 expression in addition to those with homozygous deletions

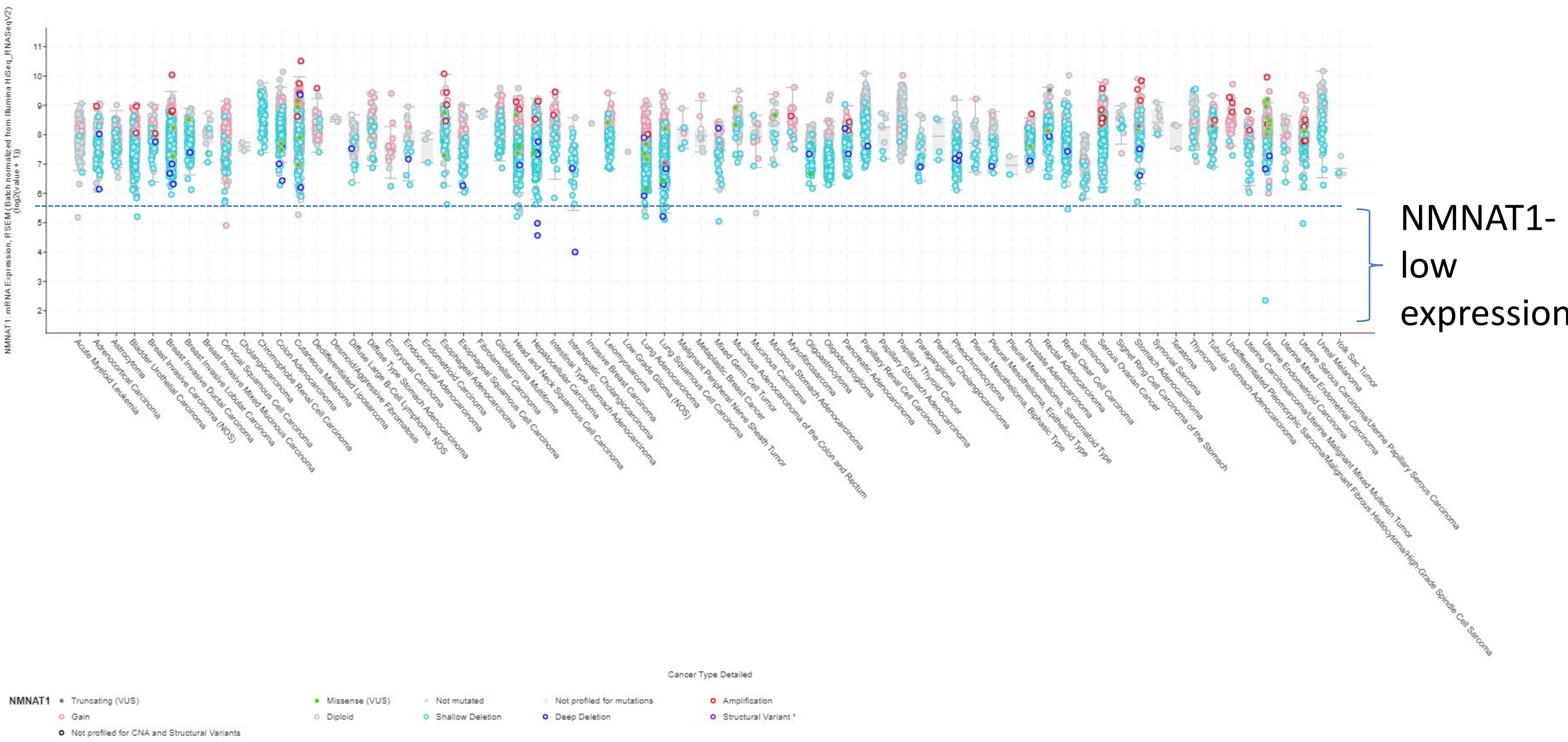

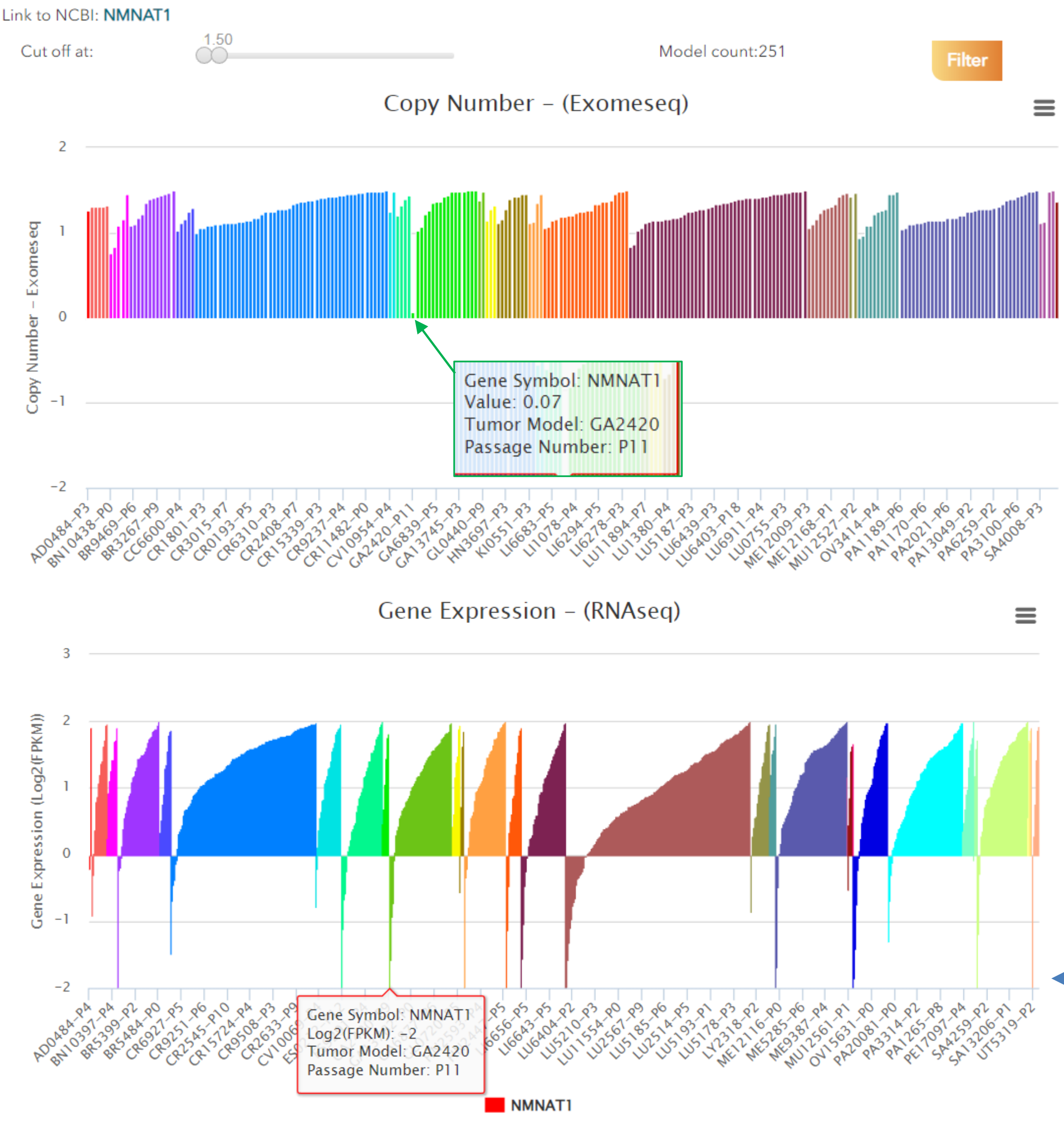


- CrownBio PDX data show Gastric Adenocarcinoma with NMNAT1 homozygous deletions
- In addition, several PDXs have near zero NMNAT1 expression

PDX lacking NMNAT1 expression
